# A mitochondrial blood-based patient stratification candidate biomarker for Parkinson’s disease

**DOI:** 10.1101/2022.02.07.479309

**Authors:** Rui Qi, Esther Sammler, Claudia P. Gonzalez-Hunt, Nicholas Pena, Jeremy P. Rouanet, Steven Goodson, Marie Fuzatti, Fabio Blandini, Kirk I. Erickson, Andrea M. Weinstein, Shalini Padmanabhan, Fox BioNet (FBN) investigators, Francesca Tonelli, Dario R. Alessi, Sruti Shiva, Laurie H. Sanders

## Abstract

Parkinson’s disease (PD) is the most common neurodegenerative movement disorder and neuroprotective interventions remain elusive. High throughput biomarkers aimed to stratify patients based on shared etiology is one critical path to the success of disease-modifying therapies in clinical trials. Mitochondrial dysfunction plays a prominent role in the pathogenesis of PD. Previously, we found brain region-specific mitochondrial DNA (mtDNA) damage accumulation in neuronal and *in vivo* PD models, as well as human PD postmortem brain tissue. In this study, to investigate mtDNA damage as a potential blood biomarker for PD, we describe a novel Mito DNA_DX_ assay that allows for the accurate real-time quantification of mtDNA damage in a 96-well platform, compatible with assessing large cohorts of patient samples. We found that levels of mtDNA damage were increased in blood derived from early-stage idiopathic PD patients or those harboring the pathogenic *LRRK2* G2019S mutation compared to age-matched healthy controls. Given that increased mtDNA damage was also found in non-manifesting *LRRK2* mutation carriers, mtDNA damage may begin to accumulate *prior* to a clinical PD diagnosis. LRRK2 kinase inhibition mitigated mtDNA damage in idiopathic PD models and patient-derived cells. The latter observations further substantiate a mechanistic role for wild-type LRRK2 kinase activity in idiopathic PD and support mtDNA damage reversal as a suitable approach to slow PD-related pathology. In light of recent advances in the field of precision medicine, the analysis of mtDNA damage as a blood-based patient stratification biomarker should be included in future clinical trials.

**One Sentence Summary:** Blood test identifies Parkinson’s patients most likely to respond to mitochondria-targeted therapeutics facilitating a precision medicine approach.

## INTRODUCTION

Parkinson’s disease (PD) is an intractable disorder characterized by progressive loss of dopaminergic neurons in the substantia nigra, resulting in tremor, rigidity, bradykinesia and postural instability. By the time a clinical diagnosis is confirmed, a significant proportion of dopaminergic neurons have degenerated (*1, 2*). Thus, neuroprotective interventions may only be successful early on during the disease course or during the prodromal phase. Even in those already diagnosed with PD, patients do not follow a uniform disease course. Rather, the heterogeneous clinical presentation of both motor and non-motor symptoms and the differing rates of disease progression support the existence of PD subtypes (*3, 4*). Thus, PD may be a group of disorders that share nigrostriatal neurodegeneration and alpha-synuclein pathology, but with distinct underlying genetic, biological and molecular abnormalities that respond differently to therapeutic approaches (*5*). An objective diagnostic test or predictive molecular marker of disease does not exist or is insufficient to identify potential subtypes thus clinical examinations remain the predominant source of PD diagnoses (*6*). The development of blood-based molecular markers to define individuals with PD that share underlying pathogenic mechanisms could transform the way clinical trials are conducted and enhance the success of disease-modifying therapies. The lack of homogenous, biomarker-defined PD cohorts has likely contributed to the failure of previous clinical trials for putative neuroprotective compounds in PD since individual differences in the underlying pathogenic mechanisms were unaccounted for. In the pursuit of neuroprotection in PD much of the focus has shifted toward precision medicine approaches, however this would require the identification of distinct pathomechanisms in individual patients (*5*). Mitochondrial function is impaired in and central to PD pathogenesis and therefore may represent one avenue to develop biomarker-validated populations in which these subsets of PD patients are amenable to putative mitochondrial targeted disease modifying therapies.

Mitochondrial dysfunction is a well-established underlying mechanism contributing to the pathogenesis of PD (*7*). Mitochondrial toxins that inhibit complex I induce pathology, nigral cell loss and parkinsonism and are used to model PD both *in vitro* and *in vivo* (*8*). Environmental exposures that target respiratory chain complexes and oxidative stress pathways are linked to an increased risk of PD in humans (*9, 10*). Moreover, patients harboring alterations in mitochondrial *polymerase gamma (POLG)* and other mitochondrial diseases exhibit nigrostriatal neurodegeneration, Lewy body pathology and clinical signs of parkinsonism (*11*). Mitochondrial genome integrity defects are observed in idiopathic preclinical models and postmortem PD brain tissues (*11, 12*). The introduction of donor mitochondrial DNA (mtDNA) from PD patients into cytoplasmic hybrid cell lines (cybrids) elicits key features of mitochondrial dysfunction and PD-related pathology, providing solid evidence that mtDNA is causal in nature for these processes (*13*). Even in monozygotic twins, the twin affected with PD demonstrates mtDNA somatic changes that match the clinical discordance (*14*). Though rare, variation in inherited pathways controlling mtDNA maintenance pathway influences PD risk (*15*).

Interestingly, both idiopathic and familial PD cases, including those with *LRRK2* mutations, are associated with mtDNA damage (*12, 16–22*). LRRK2 encodes for leucine-rich repeat kinase 2 (LRRK2), a large multidomain enzyme with tandem catalytic ROC-COR GTPase and kinase domains as well as additional motifs that are thought to contribute to protein-protein interactions (*23–25*). Pathogenic mutations in *LRRK2* are one of the most common known causes for PD, all of which cluster within its catalytic core and augment LRRK2 kinase activity with subsequent hyperphosphorylation of its endogenous RabGTPase substrates (*24, 26–28*). Multiple lines of evidence also suggest a role for LRRK2 in idiopathic PD; there is a striking resemblance between idiopathic and the genetic form due to *LRRK2* mutations, there is increased genetic risk in and around the *LRRK2* locus for idiopathic PD, and lastly one study supports non-mutation driven LRRK2 kinase activation in dopaminergic neurons in postmortem brain tissue from patients with idiopathic PD (*29–31*). Thus, LRRK2 represents a promising therapeutic target for disease modification and small molecule kinase inhibitors are currently being evaluated in clinical trials (*29, 32*).

Of particular interest is the G2019S variant, because it accounts for 4% of all familial and 1% sporadic cases worldwide, and up to 39% of cases in specific ethnic populations (*29, 33*). The penetrance of *LRRK2* G2019S is incomplete, highlighting additional genetic and environmental disease modifiers, and one such factor may be related to mtDNA dyshomeostasis (*34*). LRRK2 G2019S patient-derived fibroblasts have been associated with 7S DNA accumulation, high H-strand transcription, low mtDNA replication and mtDNA release, indicating mtDNA dysfunction (*35*). Consistent with these findings, mtDNA deletions were increased in both LRRK2 manifesting and non-manifesting PD (*36*). The role of LRRK2 in maintaining mitochondrial homeostasis is also linked to regulating damaged mtDNA release into the cytosol, at least in peripheral immune cells (*37*). Moreover, LRRK2 G2019S increases mtDNA damage in induced pluripotent stem cell (iPSC)-derived neurons from PD patients and is reversed by gene correction (*17*). In LRRK2 PD-patient derived cells, LRRK2 kinase inhibition achieved full reversal of mtDNA damage to healthy control levels; hence, mtDNA damage is a sensitive and surrogate measure of altered LRRK2 kinase activity (*16, 21*). However, even in patients with PD-linked *LRRK2* mutations, there are heterogeneous clinical and neuropathological manifestations that may reflect multiple disease pathways (*5, 38, 39*). To date, patient stratification biomarkers to help define subgroups of LRRK2 variant-carriers or idiopathic PD based on mitochondrial dysfunction do not exist, but may be warranted as a precision medicine-based approach in future clinical trials (*40*).

Previously, we found brain region-specific mtDNA damage accumulation and a parallel increase in mtDNA damage in peripheral tissues in LRRK2 and idiopathic PD patient-derived cells, models and postmortem human brain tissues (*9, 12, 16–21*). We therefore investigated for the first time, peripheral mtDNA damage as a potential blood-based biomarker in human idiopathic and LRRK2 PD. To do this, we developed the Mitochondrial DNA damage assay (Mito DNA_DX_), which allows for the accurate real-time quantification of mtDNA damage in a 96-well platform, making it compatible with evaluating large cohorts of human samples, including the Michael J Fox Foundation Fox Bionet (FBN) cohort, consisting of participants with idiopathic PD, *LRRK2-*G2019S mutation carriers with and without PD (*LRRK2-*G2019S mutation non-manifesting carriers) and age-matched healthy controls. We discovered that increased mtDNA damage is a shared phenotype in blood-derived cells from early idiopathic PD patients and those with PD carrying the *LRRK2*-G2019S mutation, compared to unaffected control subjects. Increased mtDNA damage was also found in non-manifesting *LRRK2*-G2019S mutation carriers, suggestive of accumulating mtDNA damage during the prodromal phase. Importantly, levels of mtDNA damage were not impacted acutely by PD-related medicines in a PD cohort subjected to a wash-out period. In healthy controls, test-retest reliability studies demonstrated that levels of mtDNA damage are stable over time, and alterations detected in PD samples therefore reflect the disease state. To correlate mtDNA damage with LRRK2 kinase activity, peripheral blood from FBN participants was additionally subjected to multiplexed quantitative immunoblotting for LRRK2 dependent Rab10 substrate phosphorylation as a read out for LRRK2 kinase activation status as described before (*41*). In contrast to the mtDNA damage assay, LRRK2 dependent Rab10 phosphorylation at Threonine 73 (pThr73-Rab10) did not allow discrimination between controls and those with PD and also not between those with and without the *LRRK2*-G2019S mutation. However, there was significant Rab10 dephosphorylation with *ex vivo* LRRK2 kinase inhibitor treatment compared to untreated samples for all participants, demonstrating the LRRK2 dependency of the pThr73-Rab10 phosphorylation event. LRRK2 kinase inhibition alleviated mtDNA damage in idiopathic PD models and patient-derived cells, further substantiating a mechanistic role for wild-type LRRK2 kinase activity in idiopathic PD and suggesting that reversing mtDNA damage may be beneficial to slow PD-related pathology. In light of recent advances in the field of precision medicine, PD patient stratification according to the degree of mtDNA damage may hold promise for future clinical trials.

## RESULTS

### Development of a novel PCR-based assay Mito DNA_DX_ to quantify mitochondrial DNA damage

We developed Mito DNA_DX_, which combines a high-fidelity DNA polymerase, a fluorescent dye that binds to double-stranded DNA, and optimized polymerase chain reaction (PCR) parameters that allow for the accurate real-time quantification of DNA damage in distinct loci in a 96-well platform. The principle of the assay involves amplification of a PCR fragment specific to the mitochondrial genome; less PCR product will be produced when mtDNA damage or lesions block the ability of the DNA polymerase to replicate. Thus, mtDNA damage or mtDNA repair intermediates that slow down or impair DNA polymerase progression will be detected. If equal amounts of sample DNA are amplified under identical conditions, then the quantity of PCR product can be directly compared across conditions in a statistically appropriate manner. To demonstrate the utility of Mito DNA_DX_, we first evaluated human embryonic kidney 293 (HEK293) cells incubated in hydrogen peroxide (H_2_O_2_)-containing media (an established potent and broad DNA damaging agent) for 60 minutes with a wide range of H_2_O_2_ concentrations (100-750 μM). A shift in the curve to the right in H_2_O_2_-treated HEK293 cells indicates less amplification and therefore increased mtDNA damage relative to the vehicle control (Fig. 1A). As previously observed (*42*), mtDNA lesion frequency was dependent on H_2_O_2_ concentration (Fig. 1B), and the mitochondrial amplicon is a specific PCR product (Fig. 1C). A small PCR product is used to determine mtDNA copy number (Fig. 1D). Treatment with these H_2_O_2_ concentrations for one hour was not sufficient to cause an acute loss of mtDNA copy number (Fig. 1E). The small PCR product to calculate mtDNA copy number is specific (Fig. 1F). The increase in mtDNA damage at all doses of H_2_O_2_ remains statistically significant when mtDNA lesion frequency is normalized for mtDNA copy number (Fig. 1G).

**Figure 1.**
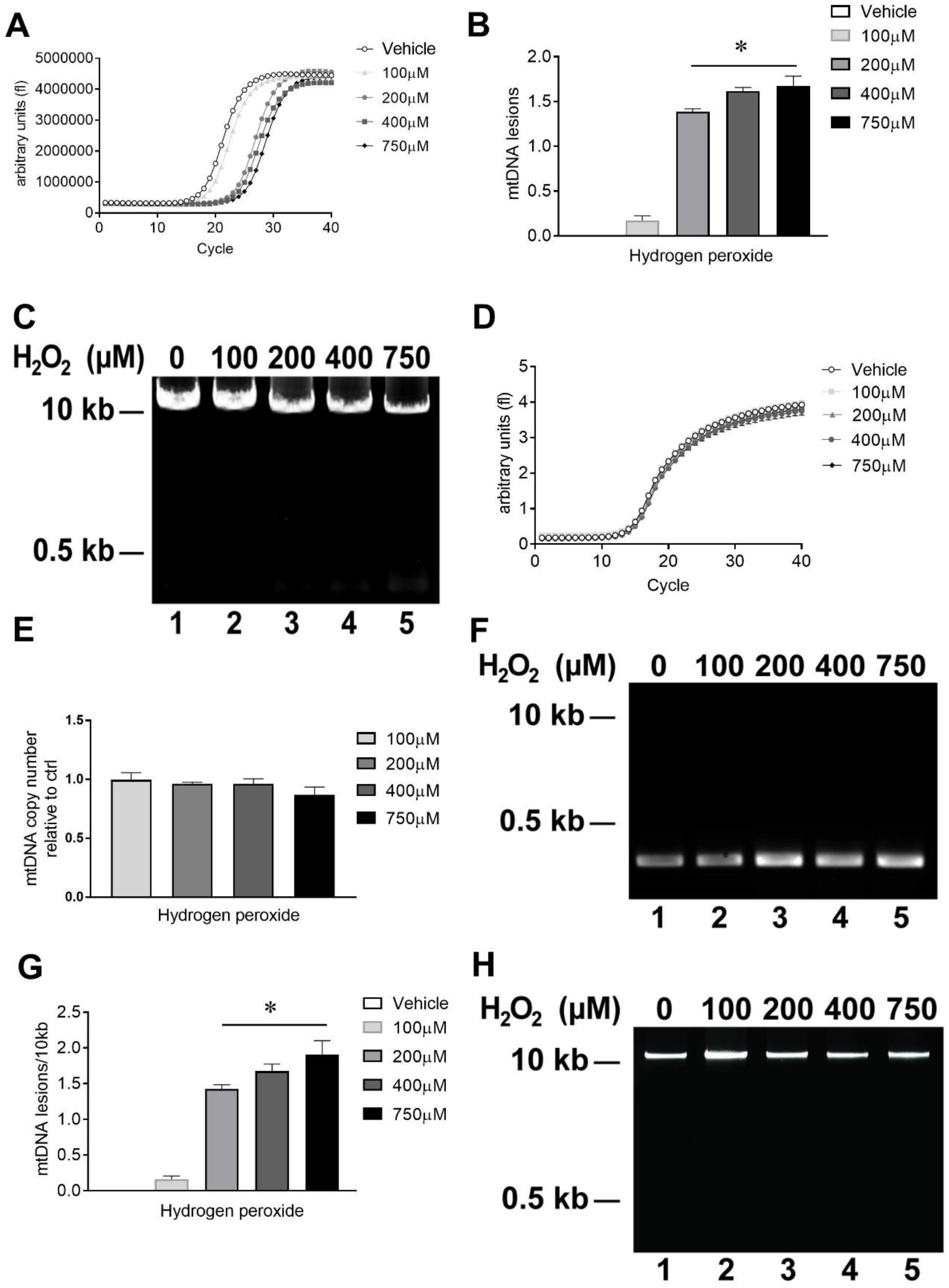
Novel Mito DNA_DX_ assay to measure mtDNA damage in real time. HEK293 cells were exposed to increasing concentrations of hydrogen peroxide (H_2_O_2_) for 1h at 37°C and total cellular DNA was isolated. **(A)** The amount of ResoLight dye fluorescence associated with each 8.9kb amplification PCR product relative to the vehicle treated control is plotted. **(B)** The decrease in relative amplification from *A* was then converted to lesion frequency using the newly developed equation as described in the methods and then applying the Poisson equation. **(C)** Representative 0.6% agarose gel depicting the specificity of the mitochondrial PCR end product. **(D)** The amount of SYBR green dye fluorescence associated with each 248bp amplification PCR product relative to the vehicle treated control is plotted. **(E)** mtDNA copy number is unchanged across conditions. **(F)** Representative 2.0% agarose gel depicting the specificity of the 248bp mitochondrial amplicon. **(G)** The mtDNA lesion frequency from *B* was then normalized both to mtDNA copy in *E* and 10kb to generate the final mtDNA lesion/10kb frequency. **(H)** Representative 2.0% agarose gel of extracted DNA using the Autogen 610L protocol that demonstrates the template DNA is of high molecular weight and quality. The Mito DNA_DX_ assay was performed in technical triplicate for each biological replicate. (*p < 0.0001, determined by one-way ANOVA with a Tukey’s post-hoc comparison). n = 3 biological replicates. Data presented as mean ± SEM.

To minimize pre-analytical variables for biomarker development and enhance throughput, the DNA extraction methodology was further optimized to be compatible with the Mito DNA_DX_ assay. The ability to amplify long (> 8kb) DNA targets is entirely dependent on template DNA integrity. However, the previously used DNA extraction methodology was time- and labor-intensive (*43*). In order to overcome these limitations, a semi-automated DNA extraction workflow utilizing an Autogen 610L, which can isolate six samples in twelve minutes, was developed to yield DNA samples that fulfill the requirements of high template quality (Fig. 1H), purity (average A260/A280 ratio = 1.71) and DNA yield (2.5 x 10^6^ cell ~ 45μg) for the Mito DNA_DX_ assay. Combining the 96-well platform with this DNA extraction methodology is more efficient and amenable to analyzing large cohorts of human samples.

### mtDNA damage observed in idiopathic PD patient-derived cells is mitigated with LRRK2 kinase inhibition

In addition to our previous findings that dopaminergic neurons selectively accumulate mtDNA damage in idiopathic PD, we found this increase in brain tissues to be paralleled by mtDNA damage in accessible peripheral tissues—such as blood—in an idiopathic PD animal model (*18*). To extend these observations to blood derived from human participants with idiopathic PD, we first compared levels of mtDNA damage in healthy control and idiopathic PD patient-derived Epstein-Barr virus (EBV)-transformed lymphoblastoid cell lines (LCL) utilizing the Mito DNA_DX_ assay (detailed demographic information in Table S1). Mitochondrial DNA damage was increased in idiopathic PD patient-derived LCLs compared to age-matched healthy controls (Fig. 2A). Mitochondrial DNA copy number was not different between idiopathic PD patient-derived LCLs compared to age-matched healthy controls (Fig. 2B).

**Figure 2.**
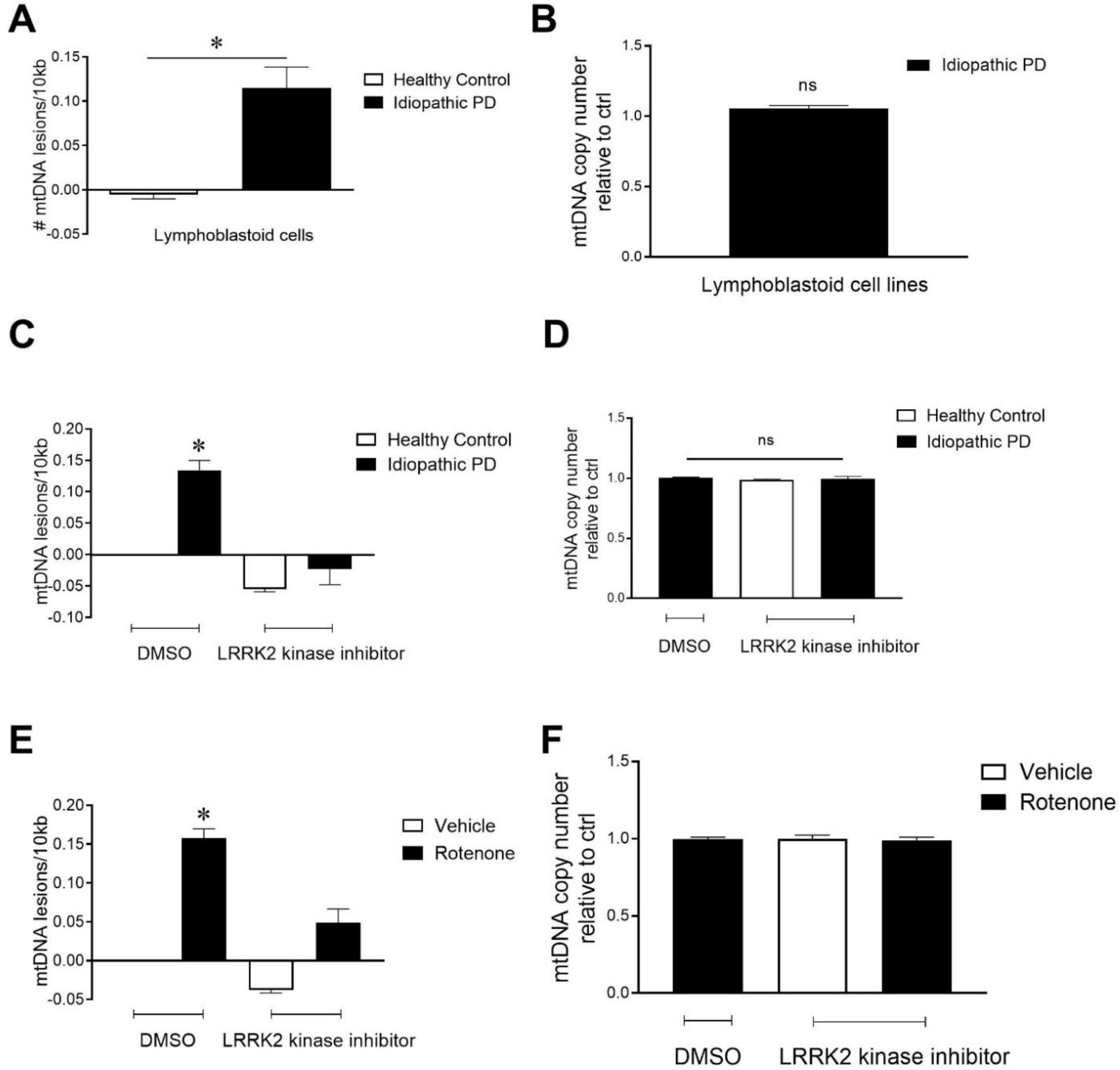
LRRK2 kinase inhibition reversed mtDNA damage in idiopathic PD patient-derived cells. **(A)** mtDNA damage was increased in idiopathic PD patient-derived LCLs (n = 4) compared to age-matched healthy controls (n = 3). (*p <0.01, determined by Student-t test). **(B)** The differences in mtDNA damage between control and idiopathic PD patient derived LCLs was not attributable to changes in steady state mtDNA levels. **(C)** Healthy control or idiopathic PD patient derived LCLs were treated with the LRRK2 kinase inhibitor, MLi-2 (1 μM for 24h). Treatment with MLi-2 reversed mtDNA damage to healthy control levels (*p<0.0001, determined by one-way ANOVA with a Tukey’s post-hoc comparison), with **(D)** no effect on mtDNA copy number. **(E)** Primary midbrain neurons exposed to rotenone (10nM) for 24h were post-treated with MLi-2 for 24h, which rescued rotenone-induced mtDNA damage (*p<0.0001, determined by one-way ANOVA with a Tukey’s post-hoc comparison). **(F)** No difference in mtDNA copy with treatment was detected. The Mito DNA_DX_ assay was performed in technical triplicate for each biological replicate. n = 3 biological replicates. ns: *p*-value >0.05. Data are presented as mean ± SEM.

Next, we tested whether culturing LCLs in the presence of a LRRK2 kinase inhibitor would affect mtDNA damage levels in idiopathic PD-derived cells. Similar to LRRK2-G2019S patient-derived LCLs (*16*), exposure of idiopathic PD patient-derived LCLs to a LRRK2 kinase inhibitor, MLi-2, restored mtDNA damage to control levels with 24 h of exposure (Fig. 2C), without a change in mtDNA copy number (Fig. 2D). Similar results were found in an *in vitro* model of idiopathic PD; primary midbrain neuronal cultures from E17 Sprague Dawley rats were treated with vehicle or a low concentration of rotenone for 24h, and then exposed to vehicle or the LRRK2 kinase inhibitor MLi-2 for 24h. After 48h of treatment, cell pellets were collected and DNA was extracted and subjected to the Mito DNA_DX_ assay. We found a statistically significant increase in mtDNA damage in primary midbrain neurons exposed to rotenone (Fig. 2E). Post-treatment with a LRRK2 kinase inhibitor reversed rotenone-induced mtDNA damage to levels of vehicle-treated primary midbrain neurons (Fig. 2E). The extent of mtDNA damage in rotenone or MLi-2 treated midbrain neurons was not related to changes in mtDNA steady state levels (Fig 2F).

### Blood-derived cells from idiopathic PD patients have elevated mtDNA damage

The next step was to evaluate whether increased mtDNA damage was also observed in non-immortalized blood-derived peripheral cells. Peripheral blood mononuclear cell (PBMC) pellets are routinely obtained for diagnostic and biomarker exploration and are currently being used for measuring target engagement and other purposes for LRRK2 and other PD-targeted clinical therapies. We first identified banked PBMCs from a previous study in which mtDNA damage could be measured from participants that included idiopathic PD and age-matched healthy controls (*44, 45*). Demographic information applicable to the samples analyzed in this cohort are summarized in Table S2; key demographic criteria were similar between PD cases and controls. Mitochondrial DNA damage levels were elevated in idiopathic PD patient-derived PBMCs compared to age-matched controls (p<0.001, Fig. 3A). Differences in mtDNA copy number were not observed between idiopathic PD and healthy controls (Fig. 3B). Mitochondrial DNA damage levels predicted PD disease status with a ROC area under the curve c-statistic of 0.83 (p<0.001, Fig. 3C). We did not detect significant correlations between mtDNA damage levels and any demographic or clinical data (Pearson or Spearman R values, all *p* values all >0.1 Table S2).

**Figure 3.**
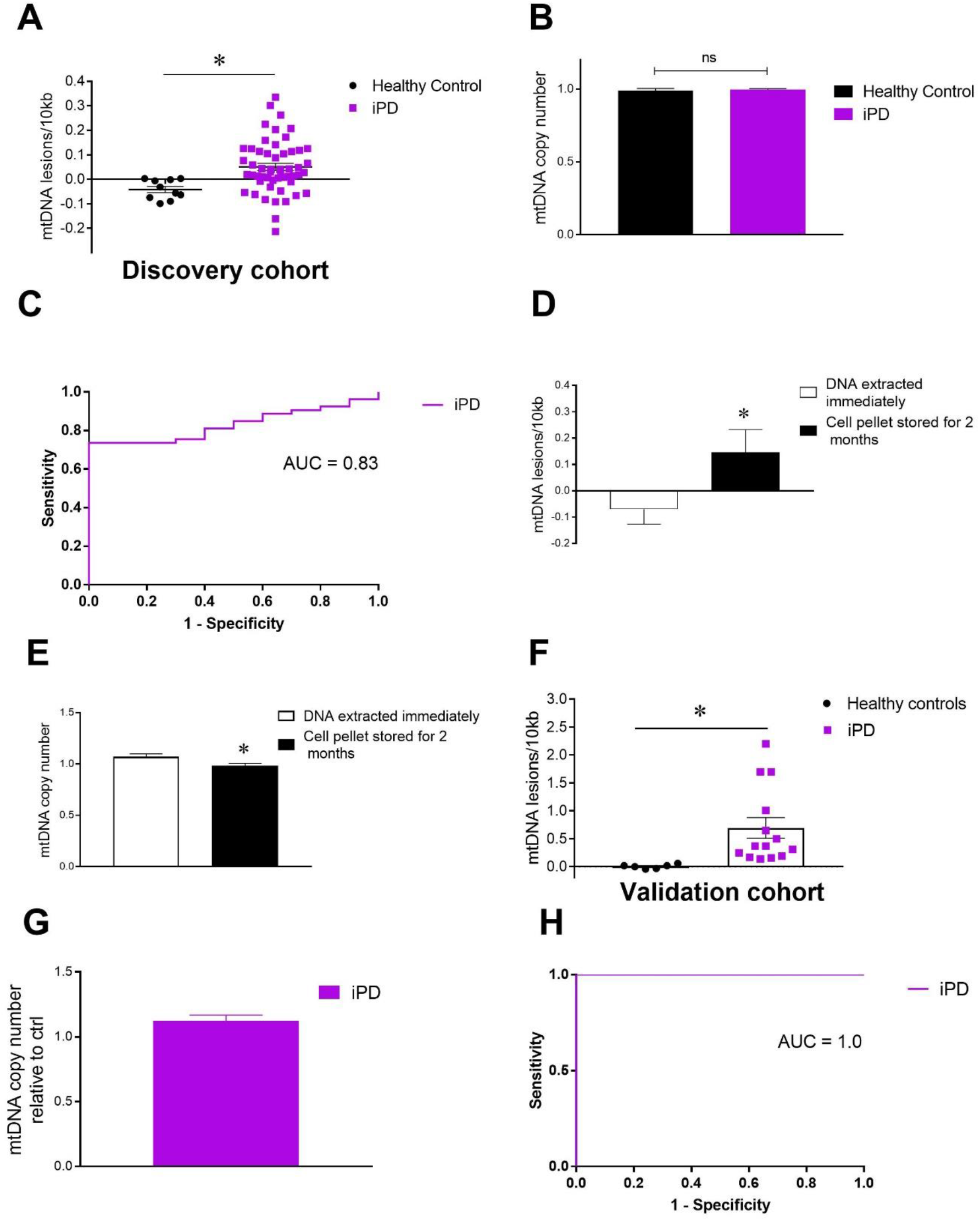
Peripheral idiopathic PD mtDNA damage levels in the discovery and validation cohort. **(A)** DNA was isolated from PBMCs derived from idiopathic PD (n=53) and age-matched healthy controls (n=10) and the Mito DNA_DX_ performed. mtDNA damage was increased in idiopathic PD compared to age-matched healthy controls (*p-value <0.001, determined by Mann-Whitney test). **(B)** The differences in mtDNA damage in healthy controls vs idiopathic PD were not attributable to changes in steady state mtDNA levels. **(C)** ROC curve showing prediction success for PD diagnosis at 83% (*p-value <0.001). **(D)** Blood samples were collected from healthy individuals and split into 2 groups - one cell pellet in which the DNA was extracted immediately. The other cell pellet sat in the freezer −20°C for two months, after which the DNA was then extracted. Storage conditions of the pellet appear to impact mtDNA damage frequency and **(E)** mtDNA copy number (*p <0.05, determined by student t-test)). **(F)** Samples were collected prospectively from idiopathic PD (n = 14) and age-matched healthy controls (n=6) and the Mito DNA_DX_ performed. mtDNA damage was increased in idiopathic PD compared to age-matched healthy controls (*p-value <0.0001, determined by Mann-Whitney test). **(G)** No differences in mtDNA copy with disease status. **(H)** ROC curve showing prediction success for PD diagnosis at 100% (p-value <0.001). The Mito DNA_DX_ assay was performed in technical triplicate for each biological replicate. ns: p-value >0.05. Data are presented as mean ± SEM. All mtDNA damage quantification were performed blinded before unblinding for clinical and demographic data.

Freezer storage duration has been reported to increase DNA damage accumulation in biological samples (*46*). Since the banked PBMC pellets from the aforementioned discovery cohort had been stored for an extensive period of time prior to DNA extraction, we explored this technical pre-analytical variable on mtDNA damage levels. To do this, healthy individuals were recruited and demographic information can be found in Table S3. From the same individual, one blood tube was used to obtain the buffy-coat derived pellet and the DNA was extracted immediately and then stored for two months at −20°C; the second blood tube was used to obtain the buffy-coat derived pellet after which the pellet was stored for two months at −20°C, after which the DNA was extracted. The two sets of samples were evaluated for mtDNA damage simultaneously. While the overall mtDNA lesion distribution was similar between the two conditions, the duration of storage of the buffy-coat derived pellet impacted baseline mtDNA lesion levels (Fig. 3D). Interestingly, mtDNA copy number was modestly affected by storage of the buffy-coat cell pellet (Fig. 3E). The Mito DNA_DX_ assay is therefore sensitive to the duration and storage conditions of the sample, which impact cell pellet integrity and ultimately shearing of the DNA, which consequently blunts the detectable mtDNA lesion range. This led us to recruit additional independent human cohorts in which we could optimally control for these pre-analytical factors.

We next determined the range of buffy-coat derived mtDNA damage levels that were collected from early idiopathic PD and age-matched healthy controls from samples collected and stored in conditions mitigating pre-analytical factors for the Mito DNA_DX_ assay. Demographics for this cohort are listed in Table S4. Buffy-coat derived samples from idiopathic early PD is associated with a statistically significant increase of mtDNA damage compared to age-matched healthy controls (p<0.0001, Fig. 3F). This increase was observed in the absence of alterations in mtDNA copy number (Fig. 3G). Mitochondrial DNA damage levels predicted PD disease status with a ROC area under the curve c-statistic of 1.0 (p<0.001, Fig. 3H). No significant correlations between any demographic or clinical data and mtDNA damage levels were observed (Pearson or Spearman R values, all *p* values all >0.1 Table S4). Overall, the magnitude and range of mtDNA damage levels are higher in the validation cohort in which the pre-analytical variables were optimized in the blood samples, compared to the discovery cohort of archived banked samples.

To address whether PD-related medicines impact mtDNA damage levels acutely, blood and clinical measures were both collected from an independent cohort of PD participants during a washout period (see methods). Early idiopathic PD and age-matched healthy controls were recruited and PBMCs assessed for mtDNA damage using the Mito DNA_DX_ assay (demographics in Table S5). Similar to the two previously described cohorts, mtDNA damage was significantly increased in PBMCs derived from early idiopathic PD patients compared to the age-matched controls (p<0.0001, Fig. 4A). Mitochondrial DNA copy number was similar across both groups (Fig. 4B). Mitochondrial DNA damage levels predicted PD disease status with a ROC area under the curve c-statistic of 0.95 (p<0.0001, Fig. 4C). Correlations for mtDNA damage levels and other demographics and clinical variables were not observed (Pearson or Spearman R values, p values all >.1 Table S5). It is noteworthy that a subset of PD subjects exhibit extremely high levels of mtDNA damage with or without a washout (Figs. 3,4). Taken together, persistent mtDNA damage is observed in circulating immune cells derived from early idiopathic PD subjects.

**Figure 4.**
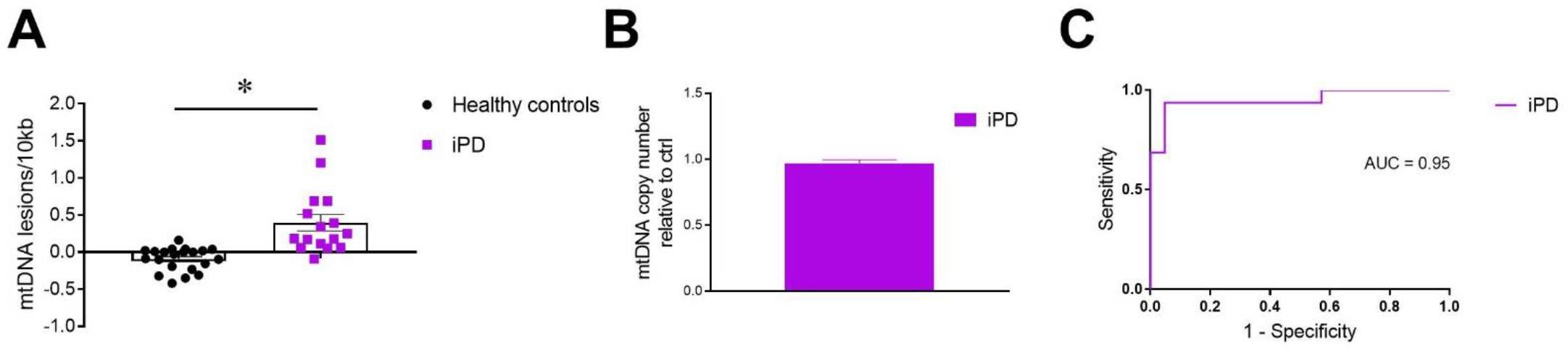
PD medications do not have an acute impact on mtDNA damage levels. **(A)** DNA was isolated from PBMCs derived from idiopathic PD (n=16) and age-matched healthy controls (n=21) and the Mito DNA_DX_ performed. Mitochondrial DNA damage was increased in idiopathic PD compared to age-matched healthy controls (*p-value <0.0001, determined by Mann-Whitney test). **(B)** The differences in mtDNA damage in healthy controls vs idiopathic PD were not attributable to changes in steady state mtDNA levels. **(C)** ROC curve showing prediction success for PD diagnosis at 95% (p <0.0001). The Mito DNA_DX_ assay was performed in technical triplicate for each biological replicate. Data are presented as mean ± SEM. All mtDNA damage quantification were performed blinded before unblinding for clinical and demographic data.

### Elevated mtDNA damage in *LRRK2* mutation carriers

Here we sought to determine whether similar increases in peripheral mtDNA damage are observed in a cohort of *LRRK2*-G2019S mutation carriers with and without a clinical diagnosis of PD, and compare to a replication independent cohort of idiopathic PD subjects. We analyzed the Fox BioNet cohort (FBN) supported by the Michael J. Fox Foundation (MJFF) which consisted of a total of 97 subjects, (n = 30 idiopathic PD, n = 28 *LRRK2-* G2019S with PD, n = 17 non-manifesting *LRRK2-*G2019S carriers and n = 22 age-matched healthy controls). The blood from the MJFF FBN cohort was collected during the clinic visit at a total of four separate academic medical centers (demographics in Table S6). Measurements using Mito DNA_DX_ in buffy-coat derived samples revealed elevated mtDNA damage in PD *LRRK2-G2019S* mutation carriers compared to non-carrier age-matched healthy controls (0.59 ± 0.1 mtDNA lesions/10kb G2019S PD versus −0.04 ± 0.05 mtDNA lesions/kb, p<0.001, Fig. 5A). Importantly, *LRRK2-G2019S* mutation non-manifesting carriers also had increased mtDNA damage relative to age-matched healthy controls (0.5 ± 0.17, p<0.01, Fig. 5A). Moreover, average levels of mtDNA damage did not differ between PD *LRRK2-G2019S* mutation carriers and *LRRK2-G2019S* non-manifesting carriers (p = 0.6). Consistent with data from previous cohorts (Figs. 3,4), mtDNA damage was increased in early idiopathic PD compared to age-matched controls (0.54 ± 0.1, p<0.001, Fig. 5A). Average levels of buffy-coat mtDNA damage derived from idiopathic PD were similar to *LRRK2-G2019S* mutation carriers—regardless of PD diagnosis (Fig. 5A, p = 0.99). Data are also presented separated by institution and demonstrate similarly robust findings across recruiting sites (Fig. S1). Mitochondrial DNA copy number was comparable across genotypes (Fig. 5B). When stratified by biological sex, female *LRRK2-G2019S* mutation non-manifesting carriers had higher mtDNA damage levels when compared to male *LRRK2-G2019S* mutation non-manifesting carriers (Fig. 5C, p<0.01). This sex difference was not observed with mtDNA copy number (Fig. 5D).

**Figure 5.**
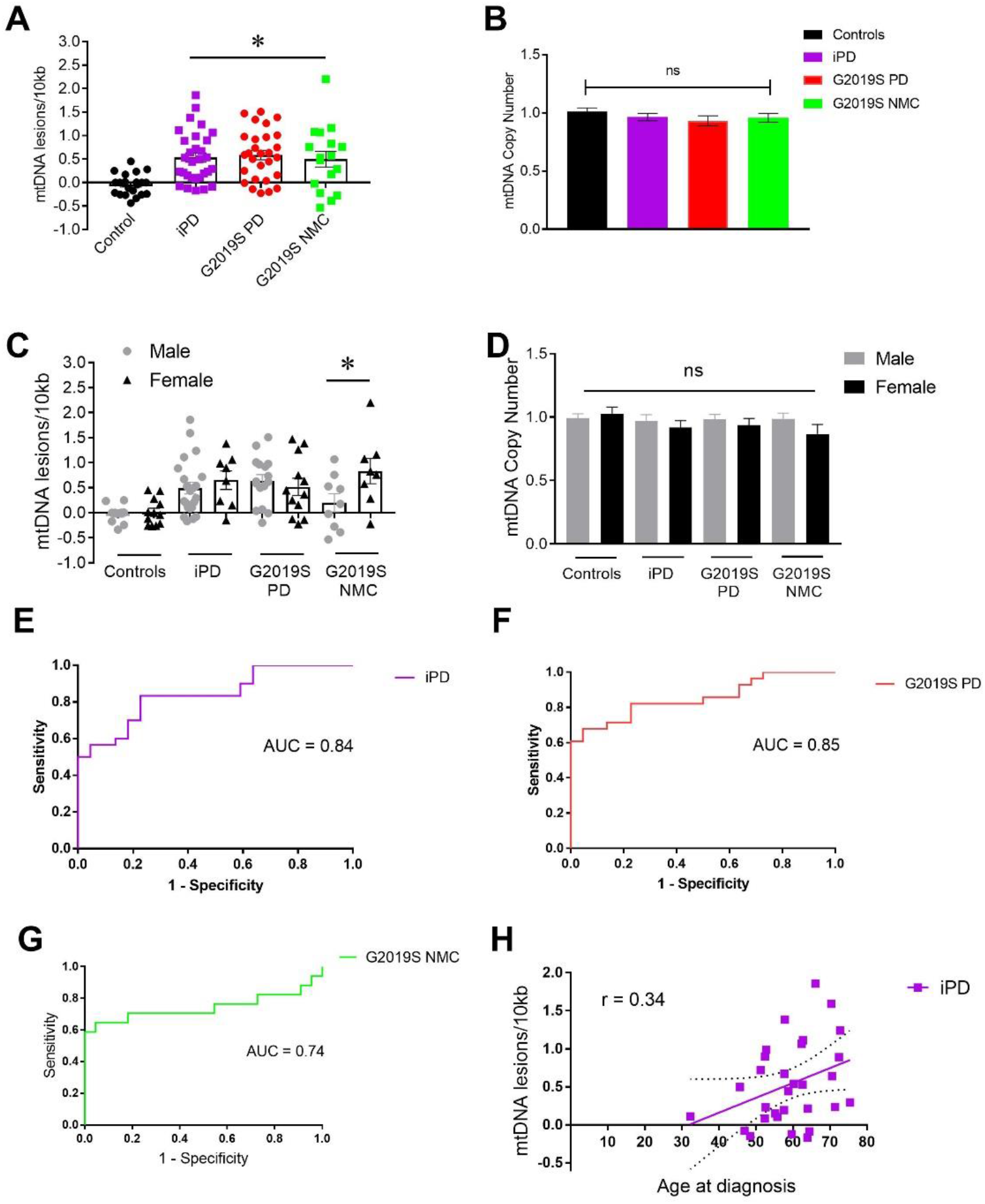
mtDNA damage is increased in both idiopathic and LRRK2 PD. **(A)** In the MJFF FBN cohort, idiopathic PD, LRRK2-G2019S PD, LRRK2-G2019S non-manifesting carriers (NMC) and age-matched healthy controls were recruited across multiple academic medical centers. Subjects with idiopathic, LRRK2-G2019S PD or LRRK2-G2019S NMC had elevated mtDNA damage relative to healthy controls (*p<0.0001, determined by one-way ANOVA with a Tukey’s post-hoc comparison). **(B)** No differences in mtDNA copy number were detected across groups. **(C)** Sex-stratification revealed that male LRRK2-G2019S NMC had lower levels of mtDNA damage compared to female LRRK2-G2019S NMCs (*p <0.01 determined by one-way ANOVA with a Sidak’s post-hoc comparison). **(D)** mtDNA copy number was similar between males and females. **(E, F)** mtDNA damage levels have excellent prediction AUC values for idiopathic and LRRK2-G2019S PD (p <0.0001). **(G)** ROC curve showing prediction success for PD diagnosis in LRRK2 G2019S NMC at 74% (p <0.01). **(H)** There is a moderate correlation between age at diagnosis and mtDNA lesion frequency (*p <0.05 pearson correlation coefficient). The Mito DNA_DX_ assay was performed in technical triplicate for each biological replicate. Data presented as mean ± SEM. All mtDNA damage quantification were performed blinded before unblinding for clinical, demographic data and *LRRK2* G2019S mutation status for the grouped analysis.

Mitochondrial DNA damage levels in idiopathic PD and PD *LRRK2-G2019S* mutation carriers strongly predicted a PD diagnosis. Mitochondrial DNA damage levels predicted a PD diagnosis in the early idiopathic PD group with a ROC area under the curve c-statistic of 0.84 (Fig. 5E, p<0.0001). Likewise, mtDNA damage levels predicted a PD diagnosis in PD *LRRK2-G2019S* mutation carriers with a ROC area under the curve c-statistic of 0.85 (p<0.0001, Fig. 5F). Perhaps not surprisingly, a lower but still acceptable ROC area under the curve c-statistic of 0.74 (p<0.01, Fig. 5G) was found for mtDNA damage levels as a predictor of a PD diagnosis in *LRRK2-G2019S* mutation non-manifesting carriers. We found a moderate positive relationship between mtDNA damage and age at diagnosis in idiopathic PD (Fig. 5H). We did not detect any other statistically significant correlations between mtDNA damage levels and demographics or clinical data (Spearman R values, all p values >0.1, Table S6).

### LRRK2 kinase dependent Rab10 Threonine 73 and LRRK2 Serine 935 biomarker phosphorylation

The LRRK2 kinase directly phosphorylates a conserved residue in the effector-binding switch-II motif of a subgroup of RabGTPases, including Rab10 at Threonine 73 (pThr73-Rab10) (*28*). In cellular models and tissues from homozygous knock-in mouse models, all *LRRK2* mutations tested so far result in hyperphosphorylation of pThr73-Rab10 mirroring LRRK2 kinase activation - with the G2019S mutation in the kinase domain resulting in an only moderate under 2-fold increase (*25, 28*). R1441 hotspot mutations in the ROC-COR GTPase domain result in a 3 to 4-fold increase in pThr73-Rab10 levels (reviewed in (*25*)). The latter has been confirmed in peripheral blood from heterozygous *LRRK2*-R1441G mutation carriers with and without PD (*47*). Phosphorylation of serine 935 of LRRK2 (pSer935-LRRK2) has historically been an important biomarker site and while it is not a direct autophosphorylation site, does not correlate with LRRK2 kinase activity, it dephosphorylates in response to Type I but not with Type II small molecule LRRK2 inhibitors and likely informs on the confirmation of the LRRK2 kinase domain (*25, 48*). We investigated pThr73-Rab10 and pSer935-LRRK2 levels in PBMCs from the MJFF FBN cohort with largely the same subjects that were analyzed for mtDNA damage (Table S6, Figure 5). For this, PBMCs were isolated from fresh peripheral blood and split into 2 equal parts for treatment with and without the specific Type I LRRK2 kinase inhibitor MLi-2 before cell lysis. Lysates were then subjected to quantitative immunoblotting multiplexed for pThr73-phosphorylated and total Rab10 as well as pSer935-phosphorylated and total LRRK2 levels, and GAPDH. Unlike for the results of the mtDNA assay, there were no statistically significant differences in pThr73-Rab10 phosphorylation levels as a readout for LRRK2 kinase pathway activation between idiopathic PD, *LRRK2-*G2019S mutation carriers, with and without PD, and age-matched healthy controls (Fig. 6A). In keeping with the LRRK2 dependency of Rab10 phosphorylation at Threonine 73, there was a significant difference between LRRK2 kinase inhibitor treated and untreated samples per group (Fig. 6A). Total Rab10 levels did not differ between the groups and also not between MLi-2 treated and untreated samples (Fig. 6B). There was also no significant difference in pThr73-Rab10 phosphorylation levels between male and female participants for each group (Fig. 6C) and no significant correlation between age at diagnosis and LRRK2 dependent pThr73-Rab10 phosphorylation levels as assessed in the idiopathic PD group (Fig. 6D). With regards to pSer935-LRRK2 phosphorylation, there was significant dephosphorylation in the samples treated with MLi-2, when compared to DMSO controls for each group (Fig. 6E). However, LRRK2 total levels were not stable with significantly lower levels in *LRRK2-*G2019S mutation non-manifesting carriers compared to idiopathic PD (p=0.0002) and generally lower in MLi-2 compared to DMSO treated samples for each group reaching statistical significance for the control (p=0.044) and idiopathic PD groups (p=0.007) (Fig. 6F). Full immunoblots blots and raw data are presented in (Fig. S2).

**Figure 6.**
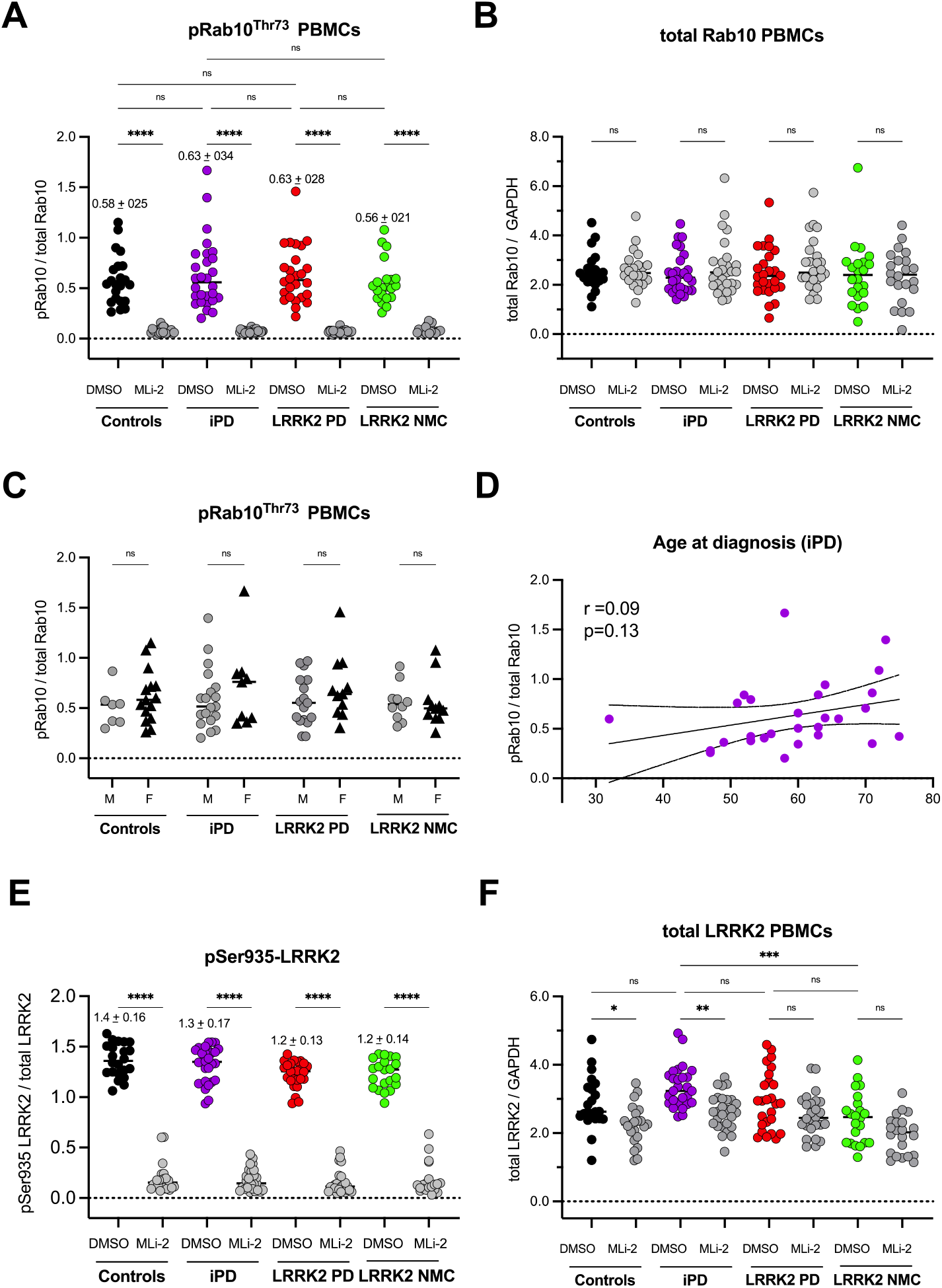
Quantitative multiplexed immunoblotting analysis of LRRK2 dependent Rab10 phosphorylation at Threonine 73 and LRRK2 Serine 935 phosphorylation in PBMCs from the MJFF FBN cohort. **(A)** LRRK2 kinase dependent phosphorylation of its endogenous substrate Rab10 at the phosphoepitope Threonine 73 (pRab10^Thr73^) was normalized against total Rab10 levels and measured in PBMC lysates from the FBN cohort that had been treated *ex vivo* with the specific LRRK2 kinase inhibitor MLi-2 (200nM, 30min) or vehicle control (DMSO). LRRK2 dependent pRab10^Thr73^ levels as a readout for LRRK2 kinase pathway activity did not show a significant difference between idiopathic PD, LRRK2-G2019S PD, LRRK2-G2019S NMC and age-matched healthy controls. There was significant dephosphorylation of pRab10^Thr7^ in MLi-2 treated samples compared to corresponding vehicle control (DMSO) treated samples for each group. **(B)** Total Rab10 levels normalized against GAPDH did not show a statistically significant difference between any of the groups or between corresponding vehicle control (DMSO) and MLi-2 treated samples. **(C)** Sex-stratification for LRRK2 dependent pRab10^Thr73^ did not reveal a significant difference between male and female sexes per group or in between groups. **(D)** There was no significant correlation between age at PD diagnosis for idiopathic PD and LRRK2 dependent pRab10^Thr73^. **(E)** Serine 935 phosphorylation of LRRK2 (pSer935 LRRK2) was measured by multiplexed quantitative immunoblotting with normalization against the total LRRK2 protein. There were no significant differences in pSer935 LRRK2 phosphorylation between the groups. There was significant dephosphorylation in LRRK2 kinase inhibitor treated samples compared to corresponding DMSO treated samples for each group. **(F)** LRRK2 total protein levels normalized against GAPDH were variable with significantly lower levels in the MLi-2 treated samples compared to DMSO for the control (p=0.044) and idiopathic PD groups (p=0.007) as well as significantly lower levels in the LRRK2-NMC group when compared to the idiopathic PD group (p=0.0002). For LRRK2 and pSer935 quantification the higher molecular weight band representing full length LRRK2 was used. Analysis was performed by one-way ANOVA with non-parametric Kruskal-Wallis multiple comparison tests except for 6C where two-way ANOVA was used. Where pairwise comparison is not indicated in the respective figures, ns: *p*-value >0.05. Data presented as mean ± SD. All immunoblotting and quantification thereof were performed blinded before unblinding for clinical, demographic data and *LRRK2* G2019S mutation status for the grouped analysis.

### Longitudinal mtDNA damage levels are stable in healthy controls

To demonstrate the internal validity of the Mito DNA_DX_ assay for PD patient assessment, test-retest reliability was measured in two ways. First, mtDNA damage levels were evaluated in buffy-coat derived samples in healthy subjects over the course of three months. Healthy individuals 50 years of age or older with no known diagnosis of neurodegenerative disease were recruited and mtDNA damage levels measured at baseline and compared to samples collected six and twelve weeks later. Of note, the broad inclusion criteria for this cohort included a range of ages, co-morbidities and medications (demographics listed in Table S7). In healthy control subjects, minimal variability in mtDNA damage levels was observed at baseline, similar to data presented in all previous cohorts (Fig. 7A). Further, optimal intra-individual stability for mtDNA damage levels was observed in these healthy subjects over twelve weeks (Fig. 7A). Intra-individual stability was also observed for mtDNA copy number over the 12-week time period (Fig. 7B).

**Figure 7.**
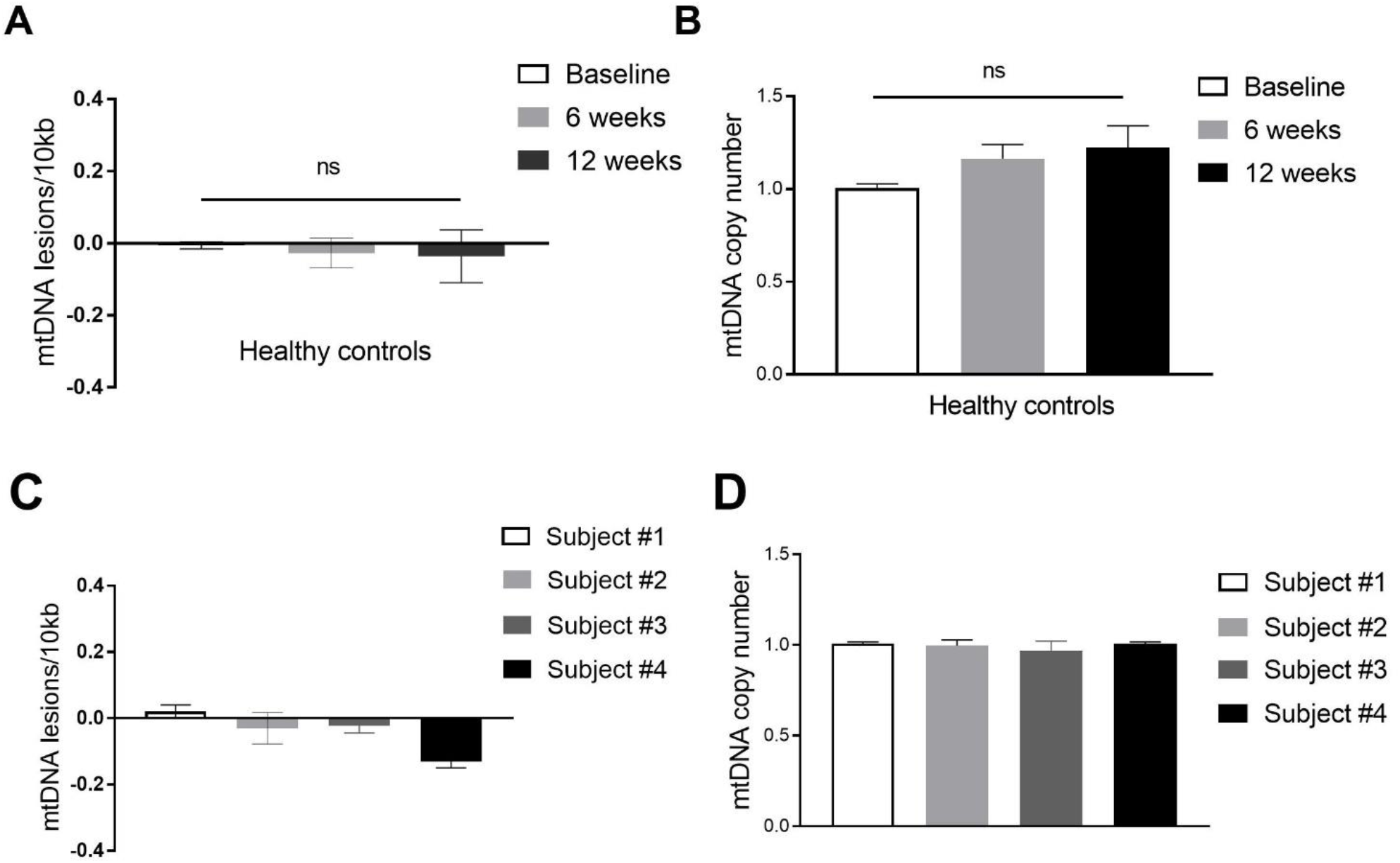
Test-retest reliability within an individual is stable over time. **(A)** Healthy subjects (without a known neurological disease) were analyzed for mtDNA damage in the same subject from a buffy-coat derived sample at three different time points (baseline, 6 and 12 weeks after the baseline was collected). mtDNA lesion frequency did not change within an individual over a 12-week period (p = 0.88). **(B)** mtDNA copy number did not statistically change with time (p = 0.15). **(C)** mtDNA damage levels were analyzed in a buffy-coat derived sample that was split approximately in half and split into two tubes. Each sample was then processed separately, and the DNA from each cell pellet compared to each other. The variability in the mtDNA lesion frequency or **(D)** mtDNA copy number was minimal in all samples analyzed. The Mito DNA_DX_ assay was performed blinded, and performed in technical triplicate for each biological replicate. ns: *p*-value >0.05. Data presented as mean ± SEM.

Next, the intra-individual variability of mtDNA damage levels was determined using a sample that was from the same individual divided into two replicates. In a subset of the healthy controls described in (Table S3), collected buffy-coat derived samples were divided equally into two separate tubes and then each cell pellet processed separately, after which then the DNA was subjected to the Mito DNA_DX_ assay. The intra-individual variability in either the mtDNA lesion frequency or mtDNA copy number was minimal in all samples analyzed (Fig. 7 C,D). Levels of mtDNA damage are exceedingly stable and consistent within and between subjects over time, indicating there is little normal fluctuation in mtDNA damage levels in the absence of overt neurodegenerative disease. Furthermore, the increased mtDNA damage observed in subjects with PD suggests loss of mtDNA maintenance associated with the disease process and not likely due to pre-analytical factors associated with sample collection, DNA extraction or the Mito DNA_DX_ assay.

## DISCUSSION

PD is a progressive neurodegenerative disease and reliable molecular biomarkers that reflect the underlying pathophysiology are urgently needed in the search for disease-modifying treatments. PD is increasingly viewed as a systemic illness that affects tissues outside of the brain and nervous system. While it is generally accepted that mitochondrial alterations in the brain play a role in the neuropathogenesis of PD, systemic mitochondrial defects have also been strongly implicated (*49*). Prior studies have shown that mitochondrial dysfunction and enzymatic deficiencies (specifically complex I inhibition) are present in the blood cells of PD patients (*8, 50, 51*). Yet, these assays require live cells and are not amenable to large scale cohorts. DNA has many characteristics that make it inherently useful for developing a biomarker, including that it is exceedingly stable and can be isolated from fresh or frozen samples. However, interrogating the mitochondrial genome poses unique challenges compared to the nuclear genome (*52*). Partly due to these technical barriers, mitochondrial DNA damage has previously not been assessed or considered as a biomarker of PD. In this study, we describe a novel PCR-based assay that measures mitochondrial genome integrity in real-time employing a medium throughput platform that produces results within 24 hours. Using Mito DNA_DX_, we identified mtDNA damage as a shared blood-based biomarker of early idiopathic and LRRK2 PD. Further, levels of mtDNA damage are responsive to LRRK2 kinase inhibition, creating the potential to identify and stratify patients with “mitochondrial PD” in the substantially large idiopathic population to test potential therapeutics in clinical trials.

Despite major advances in deciphering PD pathogenic mechanisms, drugs that are used in clinical practice to treat PD mainly work by boosting dopaminergic tone in the nigrostriatal pathway (*53*). While these therapies alleviate PD symptoms, these treatments are neither neuroprotective nor decelerate the disease. Consequently, severe disability and decrease in quality of life occur in advanced stages of the disease (*54*). To date, PD interventional clinical trials testing disease-modifying treatments have failed to show clinically meaningful effects. These failures include clinical trials testing compounds to reduce mitochondrial dysfunction—despite showing great promise in targeting mitochondrial etiology in preclinical *in vitro* and *in vivo* animal models (*55*). The factors contributing to the limited success of PD clinical trials are likely multifaceted. The interventions may be too late in the stage of disease and treatments may need to be applied earlier to enhance function or prevent further neurodegeneration. Yet, there are barriers to conducting clinical trials earlier in the disease process. One such barrier is the lack of molecular biomarkers that identify individuals at high risk for developing disease. Our findings of increased mtDNA damage in at-risk *LRRK2*-G2019S genetic carriers suggest that mtDNA damage may precede a clinical PD diagnosis and is detectable in prodromal stages. It remains to be investigated whether the non-manifesting *LRRK2* mutation carriers with elevated mtDNA damage levels develop clinical PD and could be used as a biomarker to track conversion and/or progression of the disease. Given the similarities of the distribution of mtDNA damage levels between *LRRK2* mutations carriers and early idiopathic PD, mtDNA damage may be an early marker of the disease process. Using mtDNA damage levels in at-risk populations to identify PD subjects prior to overt (and perhaps irreversible) disease opens up new avenues for earlier diagnoses, extends the window of therapeutic opportunity, and allows for faster evaluation of clinical trials outcomes.

Another consideration that may underlie the limited success of targeting mitochondria in PD clinical trials is the underappreciated complexity of mitochondrial function. Mitochondria are dynamic organelles that are critical for metabolic energy, and contribute to key cellular functions and act as gatekeepers of cell death (*56*). Mitochondrial function is tightly regulated, and when one facet is disrupted, another function may be affected, leading to mitochondrial and redox dyshomeostases. For example, antioxidant compounds such as MitoQ contain lipophilic cations and capitalize on the voltage gradient across the inner mitochondrial membrane to selectively accumulate into the mitochondrial matrix. Yet, despite the established antioxidant activity of MitoQ, unwanted effects of decreased membrane potential, mtDNA copy number and mitochondrial swelling were also observed, likely due to the high concentrations of accumulated MitoQ within mitochondria (*57, 58*). Given the dichotomy of effects on mitochondrial function, it may not be surprising that a double-blind, placebo-controlled study found no difference between MitoQ and placebo in measures of PD progression (*59*). Thus, while drugs may be beneficial in one aspect, there can be unexpected and significant mitochondrial specific toxicity, independent of the intended mechanism of action, further supporting comprehensive phenotyping of mitochondria as essential to our understanding of the complete effects of pharmacological interventions. The primary defect that initiates the cascade of mitochondrial dysfunction in PD is unknown. It is critical to better understand the mechanism of mtDNA damage accumulation and whether PD-associated mtDNA damage is upstream and causing additional mitochondrial defects or is a consequence thereof. Given that defects in mitophagy are prominent in both familial and idiopathic forms of PD (*7*), dysregulation of mitophagy could either cause or exacerbate mtDNA damage and understanding this sequence of pathogenic events and other important contributing factors will be important for future studies (*60, 61*).

Previous clinical trials have not taken into account 1) the complexity of individual differences in the underlying pathogenic mechanisms, and 2) the possibility that not all patients display molecular dysfunction in the same biological pathways (*62*). However, stratification approaches are being developed for specific subtypes of the disease (such as *α*-synuclein fibril-templating activity or mitochondrial defect subtypes) and incorporated into current trial designs (*63*). For example, coenzyme Q10, thought to improve mitochondrial function, originally failed to reach relevant clinical endpoints in a phase III randomized, placebo-controlled double-blind clinical trial; however, an ongoing trial is studying whether genetically stratified PD patients will respond preferentially (*64, 65*). A similar approach is being undertaken to investigate the potential effects of vitamin K2 in a genetically stratified cohort (*66*). Many promising compounds or treatments targeting mitochondrial impairment in PD are in the pipeline (*67, 68*). To date, a functional mitochondrial blood biomarker that could be used to establish a PD mitochondrial subtype that reflects this underlying pathophysiology has not been developed. Our data suggest that levels of mtDNA damage could form the basis of a blood-based biomarker that defines a more homogenous “mitochondrial PD” cohort. Using this method may impact future clinical trial design and represents a promising approach for more individualized treatments of PD patients.

The *LRRK2*-G2019S mutation causes mtDNA damage but homeostasis can be restored either by gene correction or LRRK2 kinase inhibition (*16, 17*). Moreover, assessment of mtDNA damage may be a measure of altered LRRK2 kinase activity levels, thereby serving as a sensitive surrogate marker (*21*). Interestingly, LRRK2 kinase activity may be elevated in idiopathic PD, even in the absence of a genetic mutation (*31*). Consistent with these findings, we found that LRRK2 kinase inhibition alleviated mtDNA damage in an idiopathic PD model and PD-patient derived cells. These results suggest a role for increased LRRK2 kinase activity in driving mtDNA damage in idiopathic PD. However, we did not find the corollary increase of Rab10 phosphorylation, a *bona fide* substrate of LRRK2, in either idiopathic PD or *LRRK2*-G2019S mutation carriers. This is consistent with previous reports for idiopathic PD and *LRRK2* G2019S mutation carriers (*69, 70*) and can be explained, at least for G2019S, with its only modest effect on LRRK2 kinase activity of under 2-fold (*25*) and possibly less in heterozygous mutation carriers. For example, LRRK2 dependent Rab10 phosphorylation is significantly increased in peripheral blood of carriers of the R1441G mutation that activates LRRK2 kinase activity 3 to 4-fold (*47*). It is possible that with methods characterized by higher sensitivity and expanded readouts for LRRK2 kinase pathway activity such as with the multiplexed targeted mass-spectrometry assay for LRRK2 phosphorylated Rabs differences may be detected (*47, 71*). We did find robust decreases in pRab10 and LRRK2 pSer935 following LRRK2 kinase inhibition with MLi-2, highlighting the value of these markers for target engagement. However, given that the LRRK2 dependent pThr73-Rab10 quantitative multiplexed immunoblotting assay deployed in this study does not distinguish between idiopathic PD, *LRRK2-G2019S* mutation carriers and healthy controls, limits its utility as a patient stratification biomarker. In the future, the same FBN samples will be subjected to our novel mass-spectrometry assay mentioned above, which will potentially allow us to further elucidate the relationship between mtDNA damage, LRRK2 kinase activity and related Rab substrate phosphorylation and improve the utility of these biomarkers.

There are several limitations of our study that could influence interpretation and translation to clinical trials. We have shown that mtDNA damage levels are increased in early idiopathic PD and in at-risk *LRRK2* genetic mutation carriers. Most critically, we do not understand how mtDNA damage phenotypes change (if at all) with the progression of the disease. In healthy controls, we found that mtDNA damage levels remain steady over a three-month evaluation period, despite broad inclusion and limited exclusion criteria for recruitment (*i.e.,* fasting was not implemented and there were no medication or co-morbidities exclusions, other than neurological criteria). While these results show that increased mtDNA damage in PD is related to the neurodegenerative disease process, how progression and severity of disease affects mtDNA damage is currently unknown. As interventional clinical trials can span years, longitudinal analyses of mtDNA damage in PD patients will be critical for interpretation of this biomarker.

A second potential limitation is that the chronic effects of PD-medicines on mtDNA damage levels cannot be fully ruled out. A wash-out experiment was conducted in which participants withheld PD-related medicines from the night before blood donation. In blood samples obtained the following morning, we demonstrated that levels of mtDNA damage levels looked similar in idiopathic PD patients with and without the wash-out, indicating that PD medicines did not acutely impact our mitochondrial measures. Testing mtDNA damage levels in drug naïve PD patients would resolve whether medicines are a significant contributor, although diagnostic accuracy is less certain in these early PD patients (*72*).

Another translational issue for developing a biomarker for mtDNA damage is that levels of damaged mitochondria in the blood may not directly reflect mitochondrial dysfunction in the central nervous system. The advantage of using blood is that the collection is minimally invasive and generally does not pose a barrier to participant recruitment for clinical trials. In the rotenone model of PD, rat blood and ventral midbrain mtDNA damage levels were both elevated, suggesting that peripheral tissues may be accurate indices of brain mitochondrial dysfunction in a preclinical model of PD (*12, 18*). Previously, we also observed accumulated mtDNA damage in vulnerable nigral neurons in postmortem PD brains (*18*). Our discovery of increased mtDNA damage in blood cells derived from idiopathic PD patients suggests that mitochondrial status in peripheral cells may mimic that of neuronal populations in the nigrostriatal pathway. Although less accessible, cerebrospinal fluid (CSF) may reflect more closely the pathology of the brain. For mitochondrial assessment, only circulating cell-free mitochondrial DNA (ccf-mtDNA) can be measured in CSF. CSF derived ccf-mtDNA is decreased in PD and levels are influenced by treatment; however, the source of this mtDNA may originate from areas not subject to neurodegeneration in PD (*73*). Nonetheless, the integrity and extent of damage of ccf-mtDNA has not been explored in PD. Lastly, though we analyzed hundreds of blood samples obtained from multiple academic centers, our findings require replication in a larger cohort, especially from more diverse populations.

In summary, we invented a novel assay to investigate mitochondrial genome integrity that can be utilized in a variety of human cellular and tissue contexts—including peripheral blood. Our data provide strong evidence for peripheral mtDNA damage in subsets of PD patients and support inclusion of mtDNA damage as a blood-based biomarker for patient enrichment in future clinical trials. These findings represent a significant advance in biomarker development that is exquisitely sensitive to the degree of mitochondrial impairment and may hold promise for PD-related interventions, such as mitochondrial enhancement therapy and other forms of precision medicine.

## MATERIALS AND METHODS

### Study Design

The objective of this study was to determine the potential of mtDNA damage as a blood-based biomarker in human PD. To do so, we developed a novel Mito DNA_DX_ assay in a 96-well platform that measures mitochondrial genome integrity. This new Mito DNA_DX_ assay was first validated using HEK293 cells treated with hydrogen peroxide. Mito DNA_DX_ was then used to assess mtDNA damage levels in four independent discovery and validation human studies in which blood was collected from PD and age-matched healthy controls. Additionally, healthy controls were recruited to obtain blood to investigate pre-analytical factors and test-retest reliability. For all human studies, sampling was approved by the local ethics committee, and all subjects signed informed consent. Additional studies examined the potential effect of LRRK2 kinase inhibition on mtDNA damage levels in PD-patient derived tissue and *in vitro* neuronal PD models, given that these classes of inhibitors are being tested as a therapeutic in clinical trials. All *in vitro* experiments were replicated at least three times. All human samples were analyzed in a blinded manner. Exclusion criteria were pre-established and exclusions (poor DNA extraction and quality, outside linear range) were made before unblinding. There was no exclusion of outliers.

### Quantifying mtDNA damage with the Mito DNA_DX_ assay

Mitochondrial DNA lesion frequency was quantified using a newly developed PCR-based assay, Mito DNA_DX_. In a 96-well MicroAmp Optical reaction plate, the reaction mixture (total 50μL) is comprised of and added in the following order: autoclaved nuclease free UV treated H_2_O to make up to 50μL volume, 15ng DNA (of picogreen verified 3ng/μL DNA (*43*) master mix (total 21.5μL) composed of 10μL 5x LongRange Buffer, 1μL BSA (100ng/mL, Sigma, A6003), 1μL dNTP mix (10mM), 2.5μL of each primer (10μM stock, 0.5μM final), 3.5μL MgCl2 (25mM), 0.5μL ResoLight (Roche) and 0.5μL KAPA Long Range HotStart DNA Polymerase (KAPABiosystems). Human primers had the following sequence: 5’-TCT AAG CCT CCT TAT TCG AGC CGA-3’ and 5’-TTT CAT CAT GCG GAG ATG TTG GAT GG-3’ for an 8.9kb mitochondrial product. Rat primers had the following sequence: 5’-GGC AAT TAA GAG TGG GAT GGA GCC AA-3’ and 5’-AAA ATC CCC GCA AAC AAT GAC CAC CC-3’ for a 13.4kb mitochondrial fragment. Each biological DNA sample was performed in technical triplicate, with both a 50% linear control and negative control (no DNA) included in each run. We used the Quant Studio3 (Applied Biosystems), and selected the 470/520 (Ex/Em) fluorescent detection filter. Cycle parameters for the human long mitochondrial amplification product was: initial denaturation at 95°C for 3 minutes, followed by 40 cycles of denaturation at 95°C for 15 seconds and extension at 66°C for 10 minutes. Cycle parameters for the rat long mitochondrial amplification product was: initial denaturation at 95°C for 3 minutes, followed by 40 cycles of denaturation at 95°C for 15 seconds and extension at 66°C for 12 minutes.

Lesion frequency was calculated using a non-linear curve fitting approach to determine the cycle number corresponding to the fluorescence of amplification for half the template using the following equation:

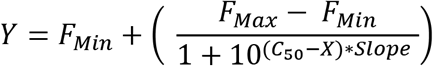

wherein Y is observed fluorescence at a selected PCR cycle, FMin is a lowest fluorescence observed during the quantitative PCR reaction, FMax is a maximum fluorescence observed during the quantitative PCR reaction, C50 is the number of cycles that produces 50% of FMax, X is the cycle number, and Slope is a slope of a curve in the linear emission range of the PCR reaction. Once the value for Y is calculated, calculations based on the Poisson equation were used, similar to those previously published (*43*).

To measure mtDNA copy number, in a 96-well MicroAmp Optical reaction plate, the reaction mixture (total 20μL) is comprised of and added in the following order: sterile UV treated H_2_O to make up to 20μL volume, 15ng DNA (of picogreen verified 3ng/μL DNA), master mix composed of 12μL which contains 10μL SYBR Green and 1.0μL of each primer (10μM stock, 0.5μM final). The human mitochondrial primers for mtDNA copy number had the following sequence: 5’-CCC CAC AAA CCC CAT TAC TAA ACC CA-3’ and 5’-TTT CAT CAT GCG GAG ATG TTG GAT GG-3’. The rat mitochondrial primers for mtDNA copy number had the following sequence: 5’-CCT CCC ATT CAT TAT CGC CGC CCT TGC-3’ and 5’-GTC TGG GTC TCC TAG TAG GTC TGG GAA-3’. The cycle parameters were initial denaturation at 50°C for 2 minutes, then 95°C for 2 minutes, followed by 40 cycles of denaturation at 95°C for 15 seconds and extension at 66°C for 1 minute. The lesion frequency for the large fragment is then normalized by dividing by the short mitochondrial amplicon as described (43). To ensure quality and specificity, PCR products were resolved on a 0.6% or 1.5% agarose gel for the larger or smaller PCR product respectively and UV light used to visualize ethidium bromide-stained gels, but were not used directly for quantification.

### Statistical analysis

GraphPad Prism (Graphpad Software, version 7 or 9) was used to perform statistical analysis, and all data are shown either as scatterplots or as means ± standard error of the mean (SEM) or standard deviation (SD). Statistical test used, number of replicates, and P value definitions are provided in the respective figure legends. Differences were considered significant when P < 0.05.

## Supporting information

Manuscript with figures

## Acknowledgments

We thank all of the patients and their families for donating blood samples, for without their generosity this study would not be possible. We are also grateful to Rajasumi Rajalingam for her administrative assistance in FBN protocol writing.

## Funding

This work was supported, in part, by grants from

The Michael J. Fox Foundation for Parkinson’s research (DRA, LHS)

Mitochondria, Aging & Metabolism Seed Grant Program (SS, LHS)

William N & Bernice Bumpus Foundation (LHS)

National Institutes of Health F31NS089111 (AMW)

Pepper Center at the University of Pittsburgh P30 AG024827 (KIE, AMW, LHS)

Medical Research Council UKRI grant MC UU 12016/2 (DRA)

Chief Scientist Office Senior Clinical Academic Fellowship SCAF/18/01 (ES)

National Institutes of Health RO1NS119528 (LHS)

## Author contributions

Conceptualization: DRA, SP, LHS,

Methodology: LHS, ES

Investigation: RQ, ES, CG, NP, SG, MF, JPR, FB, AMW, FT, LHS

Visualization: LHS, ES, FT

Funding acquisition: KIE, AMW, DRA, SS, LHS

Project administration: SS, LHS

Supervision: DRA, SS, LHS

Writing – original draft: RQ, ES, LHS

Writing – review & editing: RQ, ES, CG, NP, SG, MF, JPR, KIE, AMW, FT, DRA, SS, LHS

## Data and materials availability

All data are available in the main text or the supplementary materials. Raw data is available upon request.

