## Supplementary material for "A mitochondrial blood-based patient stratification candidate biomarker for Parkinson’s disease": Manuscript with figures

Rui Qi *et al.*

The Fox BioNet (FBN) investigators: Roy Alcalay^1^, Cheryl Waters^1^, Penelope Hogarth^2^, Tanya Simuni^3^, Danielle Smith^4^, Connie Marras^5^

^1^ Columbia University Irving Medical Center, NY, NY, USA.

^2^ Departments of Molecular & Medical Genetics and Neurology, Oregon Health & Science University, Portland, Oregon, US.

^3^ Northwestern University, Chicago, IL, USA.

^4^ Indiana University School of Medicine, Indianapolis, IN, USA.

^5^ The Edmond J Safra Program in Parkinson's disease, Toronto Western Hospital, University Health Network, University of Toronto, Canada

**Supplemental Methods**

**Semi-automated DNA extraction with the AutoGen 610L**

In general, while various kits are commercially available for DNA isolations, procedures that involve phenol/chloroform extraction must be avoided due to the introduction of artifactual DNA oxidation (*74*). High molecular weight, intact DNA is essential to efficiently amplify long genomic targets, which are required for the Mito DNA_DX_ assay. Previous protocols have used QIAGEN Genomic Tips, which was originally chosen due to the high quality DNA obtained and reproducibility from sample to sample (*74, 75*). In addition, the purified DNA is very stable, yielding comparable amplification before and after long periods of storage. The disadvantages of the QIAGEN Genomic Tip protocol are the low-throughput nature (one column at a time) and the lengthy time, three days, it takes to complete the DNA extraction procedure. The larger the target size, the higher probability that during preparation the DNA sample will have suffered a single-stranded nick or double-stranded break within the target sequence, rendering that template strand not usable for PCR amplification. As such, the comparison of mtDNA damage levels between PD and age-matched controls is highly dependent on the integrity of the DNA.

At the time of harvest, cells are trypsinized and then pelleted at 500 x g for 3 minutes. The supernatant is removed and the pellet resuspended in ice cold 1X PBS, then equal volume of buffer C1 (QIAGEN) and 3X of ice cold H_2_O. Sample is inverted to mix gently and incubate on ice for 10 minutes. The sample is pelleted at 10,000 x g at 4°C for 15 minutes and the supernatant discarded, which is critical to ensure that intact mitochondria and mitochondrial DNA are included in the sample. Samples are stored at -20°C until ready for DNA isolation. DNA was extracted with either the QuickGene DNA Whole Blood Kit L (Autogen, fk-dbl) for cells or blood, or the QuickGene DNA Tissue Kit L kit (Autogen, DT-L) for LCLs.

**Mammalian cell culture, primary rat neuronal cultures, lymphoblastoid cells lines and treatments**

HEK293 cells were maintained in DMEM/F12, GlutaMAX™ supplement (Gibco, 10565-018), Seradigm Premium Grade 10% FBS (VWR, 97068-085) and 0.5% Penicillin Streptomycin (Corning, 30-002-Cl). Cells were plated at a density of 0.3 × 10^6^ cells in a 6-well dish in triplicate for each treatment. Cells were treated at 40-50 % confluency, H_2_O_2_ (30%, Sigma) was serially diluted and added to media to achieve a final concentration in the range of 100-750μM and treated for 1h. At the time of harvest, cells were collected and DNA extracted as described above. Idiopathic PD patient (n = 4) and healthy subject control (n = 3)-derived lymphoblastoid cell lines (LCL) were obtained from the NINDS Coriell biorepository (ID numbers are indicated in Table S1). There was no statistical difference in age between the idiopathic PD patient and healthy control subjects (P = 0.92). LCLs were cultured at 37°C, 5% CO_2,_ in RPMI-1640 (Sigma-Aldrich, R8758), 15% heat-inactivated fetal bovine serum (VWR Seradigm, 97068-091) and 0.5% Penicillin/Streptomycin (Corning, 30-002-CI). Cells were passaged every 3-4 days, and passage number did not exceed 20. Primary ventral midbrain neurons were prepared following previously published protocols (*16*). Primary ventral midbrain neuronal cultures were treated with either DMSO or 10nM rotenone for 24h on DIV 7. Neurons were then treated with MLi-2 for 24h and DNA was collected on DIV 9.

**Participants**

*Discovery cohort subjects (Fig. 3A-C).* PD Patients attending the Center for Parkinson’s disease and Movement disorders of the IRCCS National Neurological Institute C. Mondino, Pavia, Italy were recruited for the study. The protocol was approved by the Ethical Committee of the IRCCS National Neurological Institute C. Mondino and all subjects received a comprehensive description of the study and signed an informed consent before being enrolled (*44, 45*). Healthy subjects matched for age and sex distribution without history of degenerative or cerebrovascular diseases, and without cognitive impairment or other neurologic disorders were recruited as normal controls. None of the subjects participating in the study was suffering from inflammatory or infectious diseases or was taking drugs capable of interfering with white blood cells (*44*).
Patients were staged according to the Hoehn and Yahr criteria and evaluated with the Unified Parkinson’ Disease Rating Scale (UPDRS subscale III motor) at the time of the enrollment. Clinical evaluation was achieved at the visit when patients also underwent a venous blood sampling (30 mL) in the morning after overnight fast. Maximal caution was achieved so that sampling was performed 2-3 hours from the last L-DOPA intake. Blood was collected from the antecubital vein into vacuum tubes containing ethylenediamietetraacetic acid (EDTA), as anticoagulant (Terumo Europe, Belgium). The blood was diluted to 50 mL with phosphate buffered saline (PBS), pH 7.4, layered over half of its volume of Lympholite (Sigma) and centrifuged at 450 x g for 20 minutes. After removing the upper layer, the PBMC band at the interface was transferred into a new tube, washed with PBS and centrifuged again. The cell pellet was resuspended in 5 mL of PBS and cells counted using a Scepter 2.0 Handheld automated cell counter (Millipore). Samples were split in aliquots containing 2 x 10^6^ PBMCs and stored at -80°C for future evaluation. Inclusion criteria was age 30 or older (at diagnosis if applicable), ability to provide consent, healthy controls had no resting tremor, bradykinesia or any other signs of atypical parkinsonism, and exclusion criteria included being a participant in a clinical trial in the last 6 months, treatment for cancer in the last 5 years, or a family history of PD. All quantitative mtDNA damage analysis was performed blinded for clinical and demographic data.

*Validation cohort subjects (Fig. 3F-H).* Participants were recruited from the University of Pittsburgh Movement Disorders Clinic registry for the parent observational study examining the association of physical activity and cognitive performance in PD (*76*). Eligibility criteria for all participants included age between 50 and 80 years, fluent in English, normal or corrected to normal sensory ability; a clinical diagnosis of idiopathic PD, Hoehn & Yahr score of <2, and a stable medication regimen for at least 1 month. Exclusion criteria for all participants included self-reported neurological or current psychiatric conditions (other than PD), significant cardiovascular disease or recent cardiac event, hospitalization in the past 3 months, score <22 on the Montreal Cognitive Assessment (MoCA), and magnetic resonance imaging contraindications. For the PD participants, exclusion criteria also included atypical Parkinsonism and current participation in an intervention trial. Health information was assessed by self-report and not by medical record review. All participants were approached to participate in the ancillary blood draw for mtDNA damage assessment and 15 of the PD participants and all of the control participants agreed to the blood draw. PD patients completed assessments while in the “ON” phase of their medication schedules (i.e., within 1–2 hours of ingesting medications), such that assessments reflected participants’ medication-adjusted performance. Participant demographic data are shown in Supplemental Table 4. Venous blood was collected into a sodium citrate tube (BD Bioscience) and buffy-coat preparations isolated. 8mL of blood was collected and within 2 hours after samples were collected, the blood sample(s) was centrifuged at 1500 x g at room temperature for 20 minutes with the brake turned off. Plasma was removed and the buffy-coat transferred to a 15mL conical tube; the volume was adjusted to 2mL with 1X PBS. 1 mL ice-cold Buffer C1 (QIAGEN) and 3 mL of ice-cold distilled autoclaved water was added and then the sample inverted several times to mix and incubated on ice for 10 minutes. Samples were centrifuged at 4°C for 15 minutes at 10,000 x g. The supernatant was removed and the pellet was stored at -20°C prior to DNA isolation. Both the parent and ancillary blood draw was approved by the University of Pittsburgh IRB and all participants provided informed consent. All quantitative mtDNA damage analysis was performed blinded for clinical and demographic data.

*Validation cohort with idiopathic PD subjects with a wash-out (Fig. 4A-C).*

Participants were recruited from the University of Pittsburgh Movement Disorders Clinic registry. Eligibility criteria for all participants included age greater than 50 years old, fluent in English and for PD participants a clinical diagnosis of idiopathic PD. Exclusion criteria for all participants included self-reported neurological or current psychiatric conditions (other than PD), and participation in an intervention clinical trial within the last six months. Health and demographic information was assessed by a medical health questionnaire and by medical record review. PD participants delayed PD-related medicines for at least 12 hours prior to the visit and withheld medication until blood was collected and testing was completed. PD patients completed assessments while in the “OFF” phase of their medication. PD participants then resumed their normal medication schedule, and the UPDRS was then assessed while on the “ON” phase of their medication, (i.e., typically within an hour of ingesting medications). Participant demographic data are shown in Supplemental Table 5. Venous blood was collected into a sodium citrate tube (BD Bioscience) and buffy-coat preparations isolated. In brief, 8mL of blood was collected and within 2 hours after samples were collected, the blood sample(s) was centrifuged at 1500 x g at room temperature for 20 minutes with the brake turned off. Plasma was removed and the buffy-coat transferred to a 15mL conical tube; the volume was adjusted to 2mL with 1X PBS. 1 mL ice-cold Buffer C1 (QIAGEN) and 3 mL of ice-cold distilled autoclaved water was added and then the sample inverted several times to mix and incubated on ice for 10 minutes. Samples were centrifuged at 4°C for 15 minutes at 10,000 x g. The supernatant was removed and the pellet was stored at -20°C prior to DNA isolation. The study was approved by the University of Pittsburgh IRB and all participants provided informed consent. All quantitative mtDNA damage analysis was performed blinded for clinical and demographic data.

*Michael J. Fox Foundation Fox Bionet (MJFF FBN) Cohort (Fig. 5) and blood sample collection for analysis of mtDNA damage and LRRK2 dependent phosphorylation.*

The MJFF FBN cohort consisted of 111 participants recruited from 4 participating US movement disorder centers (Oregon Health & Science Center, Columbia University Medical Center, University of Pennsylvania, Northwestern University) between 2019-2020 and made up of 4 groups including healthy controls (n=31), idiopathic PD (n=30) and carriers of the *LRRK2* G2019S mutation (n=50) with Parkinson’s (n=28) and non-manifesting carriers (n=22) as well as healthy controls (n=22). 87 of the 111 subjects took part in both studies, the mtDNA damage assay as well as the LRRK2 dependent Rab10 phosphorylation assay, 14 participants only participated in the former and 10 only in the latter. Demographics including sex and age as well as age at PD diagnosis and disease duration and can be found in Supplemental Table 6. PD diagnosis was defined according to the MDS clinical diagnostic criteria for Parkinson's disease (*77*). Severity of motor symptoms and the presence of motor complications was assessed using part III and IV of the Movement Disorder Society—Unified Parkinson’s disease rating scale (MDS-UPDRS-III, -IV) and Hoehn and Yahr stage (*78*). The study was approved by the respective local ethics committees and all participants gave written informed consent.

Two samples of venous blood were collected; one into a sodium citrate tube (BD Bioscience) for buffy-coat preparations for mtDNA damage analysis as described above in the validation cohort and the pellet stored at -20 ^o^C until shipment to Duke University for analysis. The other blood sample was collected into an Ethylenediaminetetraacetic acid (EDTA) tube for peripheral blood mononuclear cells (PBMCs) isolation with subsequent treatment with and without the specific LRRK2 kinase inhibitor MLi-2 as reported before (*41*). Briefly, PBMCs were isolated from 15 ml of fresh blood via density centrifugation using Ficoll-Paque PREMIUM (GE Healthcare, Cat# 17-5442-02) and the SepMate™ tube (STEMCELL, Cat# 85450, 50 ml capacity). PBMCs were then washed in phosphate buffered saline (PBS) and pelleted for resuspension in 10.5ml of PBS supplemented with 2% fetal bovine serum (FBS). The PBMC suspension was then divided into 2 aliquots for treatment with 200nM MLi-2 LRRK2 kinase inhibitor or equivalent volume of DMSO for 30 minutes. PBMCs were then pelleted by centrifugation, the supernatant was discarded, and each cell pellet was lysed in 100ul of Di-isopropylfluorophosphate (DIFP) containing lysis buffer. Lysates were then snap frozen and stored at -80 ^o^C until shipment to the University of Dundee for analysis. All quantitative mtDNA damage and western blot analysis was performed blinded for clinical and demographic data as well as *LRRK2* G2019S mutation status.

*Healthy control cohorts (Fig. 3,7).* Participants were recruited from the Duke University healthy volunteer registry at the Duke Early Phase Research Unit (DEPRU) or from caregivers of PD patients seen at the Movement Disorders Clinic. Eligibility criteria for all participants included aged 50 years and older, able to read and speak English, not a current smoker, and no cancer treatment in the last five years. Exclusion criteria for all participants included lack of a known neurological degenerative disease or participation in an interventional clinical trial within the last six months. Participant demographic data are shown in Supplemental Tables 3 and 7. Venous blood was collected into a sodium citrate tube (BD Bioscience) and buffy-coat preparations isolated. Samples were collected as described above in the validation cohort with the following modification: for data presented in Fig. 3, two blood samples were collected from each individual. One sample was pelleted and the DNA extracted immediately. The other sample was pelleted and stored in the freezer for two months at -20°C and then the DNA subsequently isolated. For data presented in Fig. 7 A,B blood was collected at the initial visit, six weeks and 12 weeks after the initial sample collection. In a subset of subjects, (Fig. 7 C,D) the buffy-coat derived sample was evenly divided into two tubes and processed separately. This study was approved by the Duke University IRB and all participants provided informed consent. All quantitative mtDNA damage analysis was performed blinded for treatment.

**LI-COR® quantitative immunoblotting**

*Quantitative multiplexed immunoblotting analysis of MJFF FBN cohort samples*: All corresponding LRRK2 kinase inhibitor and DMSO vehicle control treated PBMC lysates were cleared by centrifugation at 20 800g for 10 min at 4 °C. Supernatant cell lysates were collected, protein concentration was quantified by the Pierce BCA Protein assay (Pierce, Therma Fisher, Cat# 23225) and lysates were mixed with 4 × SDS–PAGE loading buffer [250 mM Tris–HCl, pH 6.8, 8% (w/v) SDS, 40% (v/v) glycerol, 0.02% (w/v) Bromophenol Blue and 4% (v/v) 2-mercaptoethanol] to a final total protein concentration of 1 µg/µl and heated at 70 °C for 10 min. All quantitative immunoblotting was performed blinded for clinical and demographic data as well as *LRRK2* G2019S mutation status. 10 µg per sample and analysis was loaded in duplicates onto a NuPAGE 4–12% Bis–Tris Midi Gel (Thermo Fisher Scientific, Cat# WG1403BOX) and electrophoresed at 130 V for 2 h with the NuPAGE MOPS SDS running buffer (Thermo Fisher Scientific, Cat# NP0001-02). At the end of electrophoresis, proteins were electrophoretically transferred onto a nitrocellulose membrane (GE Healthcare, Amersham Protran Supported 0.45 µm NC) at 100 V for 90 min on ice in transfer buffer (48 mM Tris–HCl and 39 mM glycine). Transferred membranes were blocked with 5% (w/v) skim milk powder dissolved in TBS-T [20 mM Tris–HCl, pH 7.5, 150 mM NaCl and 0.1% (v/v) Tween 20] at room temperature for 1 h. The membranes were then cropped into three pieces, namely the ‘top piece’ (from the top of the membrane to 75 kDa), the ‘middle piece’ (between 75 and 30 kDa) and the ‘bottom piece’ (from 30 kDa to the bottom of the membrane). The top pieces were incubated with rabbit monoclonal anti-LRRK2 pS935 antibody (UDD2, MRC PPU Reagents and Services, used at 1:1000 dilution) multiplexed with the mouse monoclonal anti-LRRK2 C-terminus total antibody (75-2553, Antibodies Inc./NeuroMab, 1:1000) diluted in 5% (w/v) skim milk powder in TBS-T to a final concentration of 1 µg/ml for each of the antibody. The middle pieces were incubated with mouse anti-GAPDH antibody (sc-32233, Santa Cruz Biotechnology, 1:10.000) diluted in 5% (w/v) skim milk powder in TBS-T to a final concentration of 50 ng/ml. The bottom pieces were incubated with rabbit monoclonal MJFF-pRAB10 antibody (ab230261, Abcam, 1:1000) multiplexed with mouse MJFF-total Rab10 monoclonal antibody (0680–100/Rab10-605B11, Nanotools, 1:1000) diluted in 2% (w/v) bovine serum albumin in TBS-T to a final concentration of 0.5 µg/ml for each of the antibody. All membranes were incubated in primary antibody overnight at 4 °C. Prior to secondary antibody incubation, membranes were washed three times with TBS-T for 10 min each. The top and bottom pieces were incubated with goat anti-mouse IRDye 680LT (#926-68020) secondary antibody multiplexed with goat anti-rabbit IRDye 800CW (926-32211, LI-COR) secondary antibody diluted in TBS-T (1: 10,000 dilution) for 1 h at room temperature. The middle pieces were incubated with goat anti-mouse IRDye 800CW (926-32210, LI-COR) secondary antibody diluted in TBS-T (1: 10,000 dilution) at room temperature for 1 h. Membranes were then washed in TBS-T three times for a duration of 10 min each. Membranes were then scanned using the LI-COR Odyssey CLx imaging system.

*Quantification of the LI-COR immunoblot analysis*. Quantification of the protein bands was performed on the scanned images using the Odyssey Scan band tool in a blinded manner with regards to genotype and clinical status of the participants using the Image Studio software by LI-COR. Each sample set including DMSO and MLi-2 treated PMBC lysates for each participant was run in duplicates and two independent replicate immunoblotting experiments were performed and used for quantification. The intensities of the total LRRK2 and total Rab10 protein bands were normalized to that of the housekeeping protein GAPDH (total LRRK2/GAPDH and total Rab10/GAPDH) while the specific posttranslational phosphorylation modifications were quantified against the multiplexed total target protein irrespective of modification (Thr73-pRab10/total Rab10 and Ser935-pLRRK2/total LRRK2). Inter-gel variability was controlled for by normalization against the same control sample on each gel per set of experiments. Once LI-COR quantification was completed, participant information was unblinded for statistical analysis.

**Supplementary Figures**

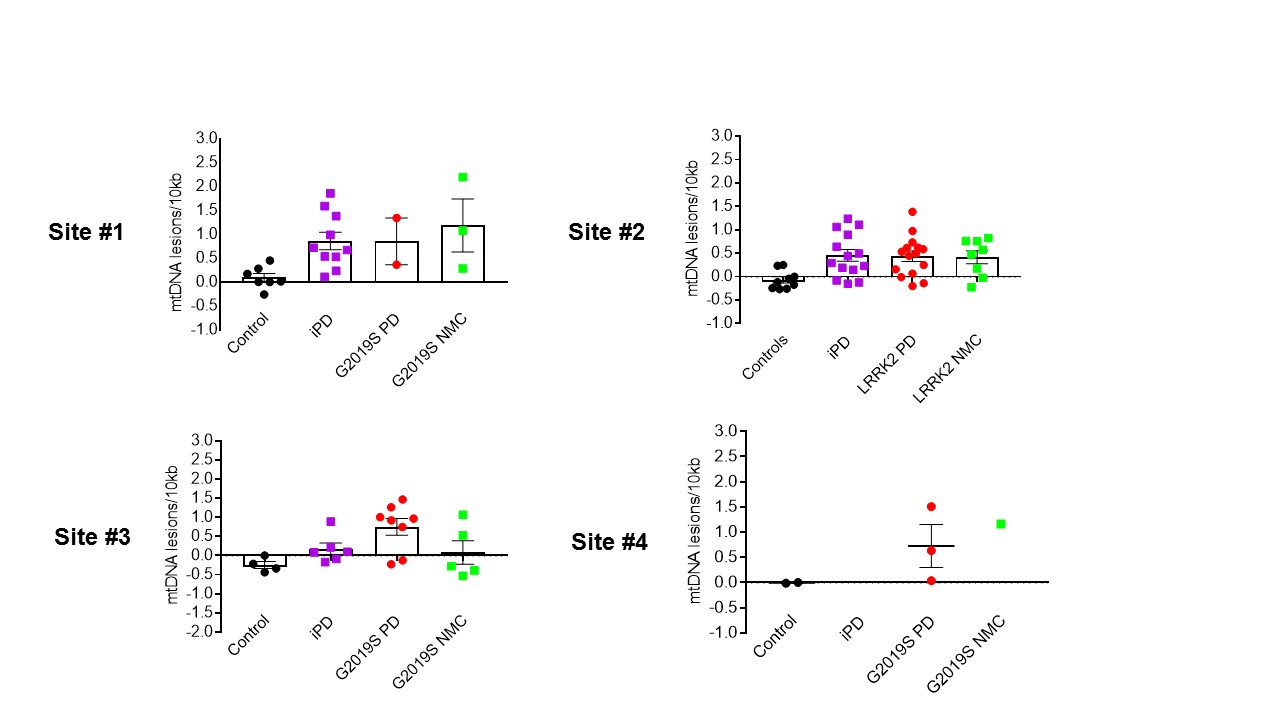

**Supplemental Figure 1. mtDNA damage levels in human participants across individual recruiting academic sites.** DNA was extracted from buffy-coat either from idiopathic PD, LRRK2-G2019S PD or LRRK2-G2019S non-manifesting carriers (NMC) and healthy controls. Mitochondrial DNA damage was analyzed using Mito DNA_DX_ in a blinded fashion. After the analysis was performed data was separated by each recruiting site. Similar increases in mtDNA damage were robustly observed in idiopathic or LRRK2 PD and LRRK2 G2019S NMC relative to healthy controls independent of academic recruiting site.

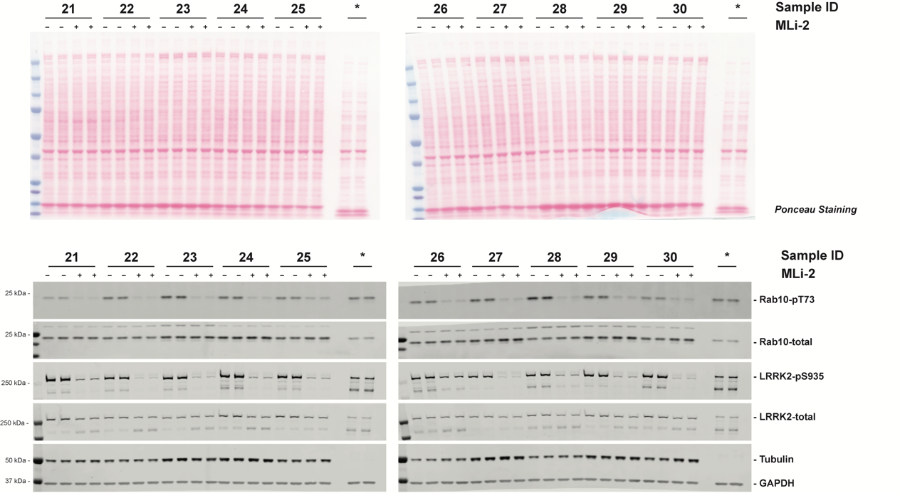

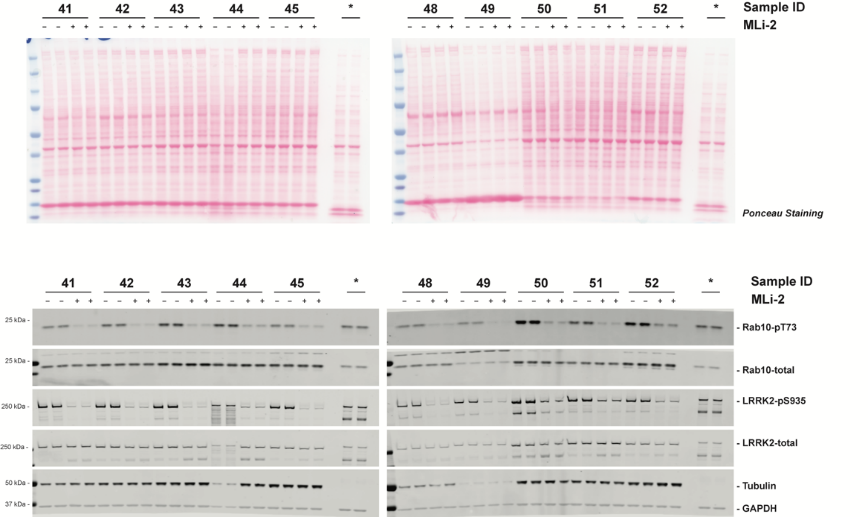

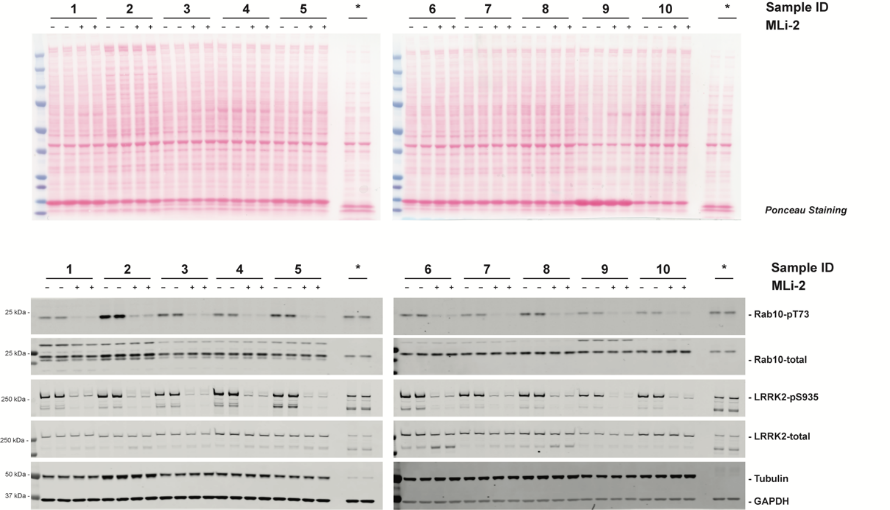

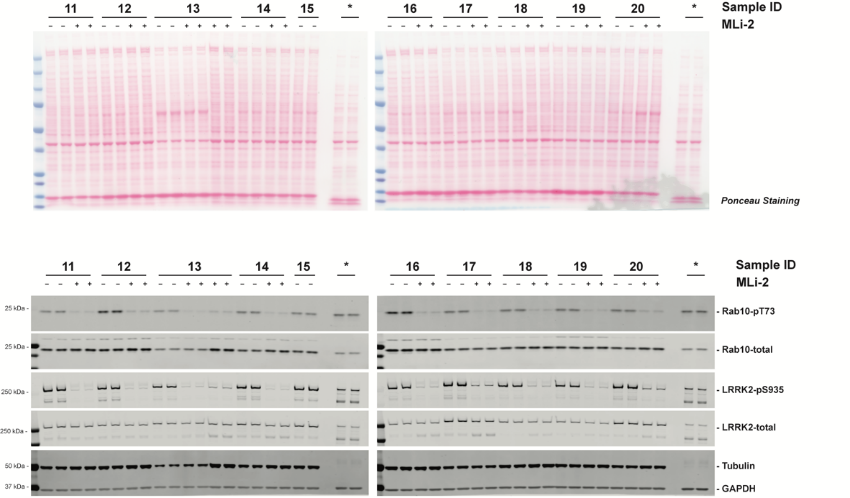

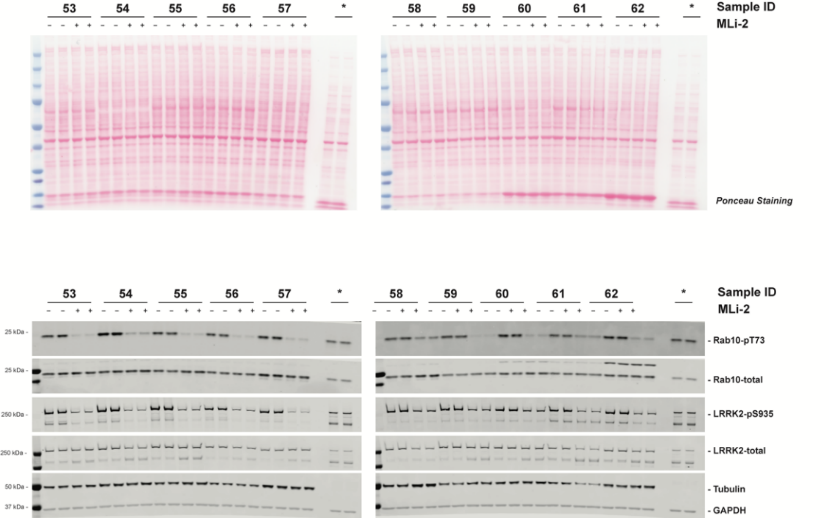

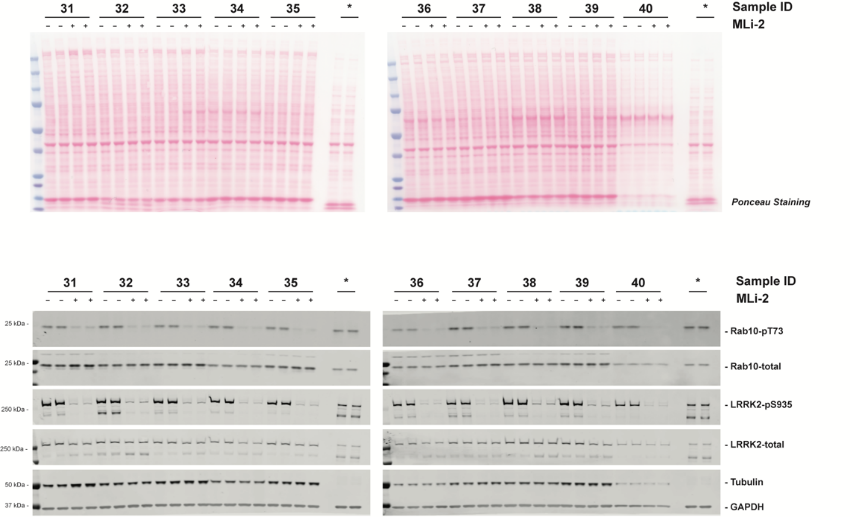

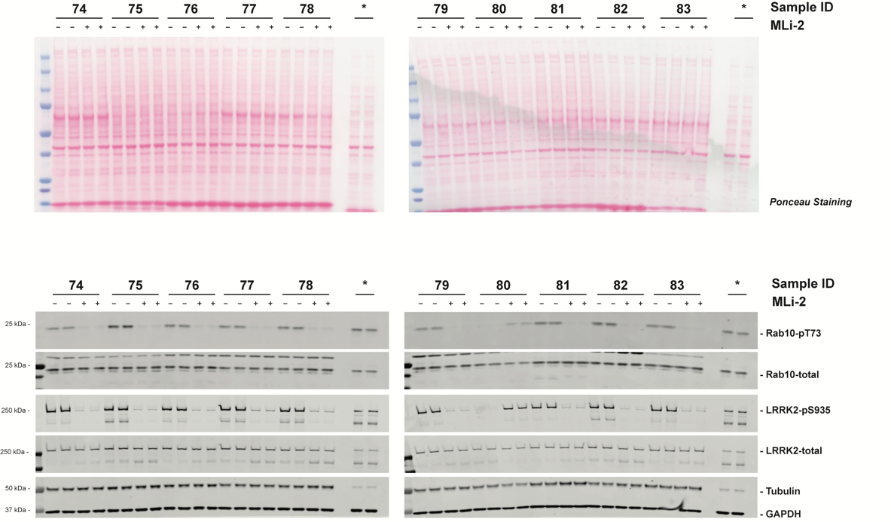

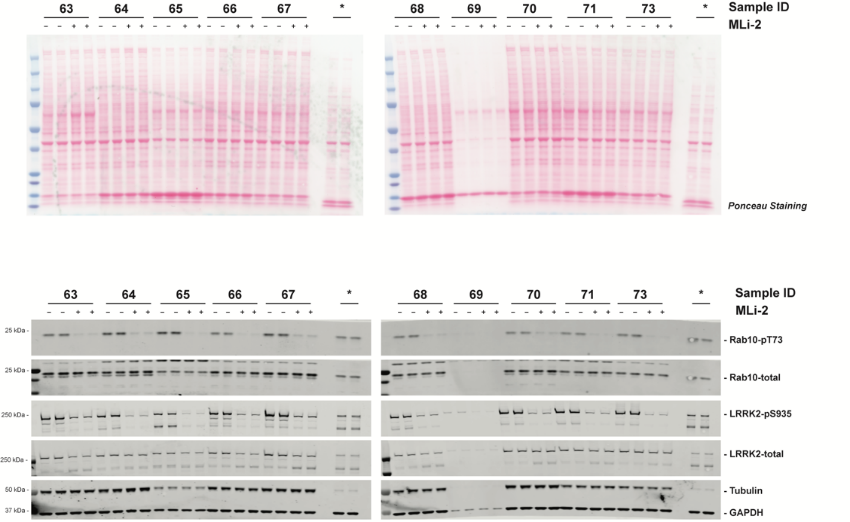

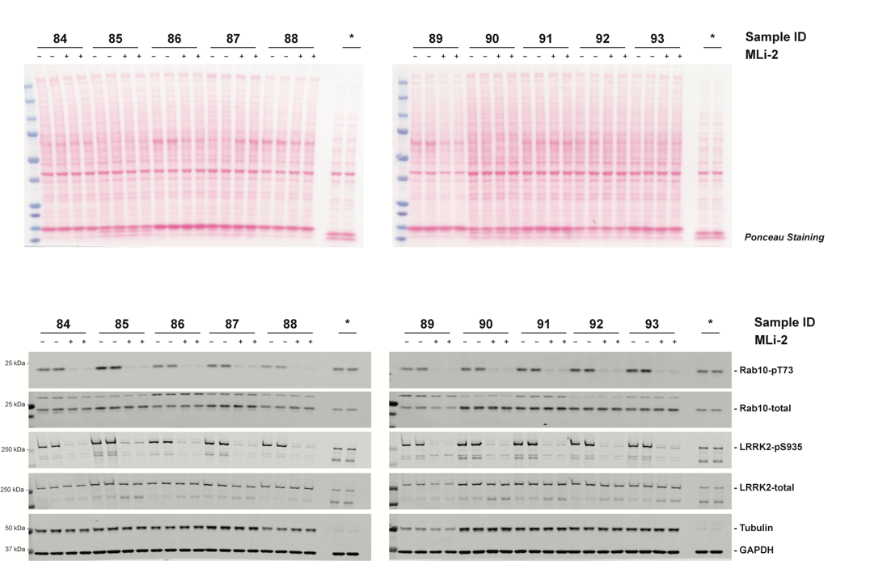

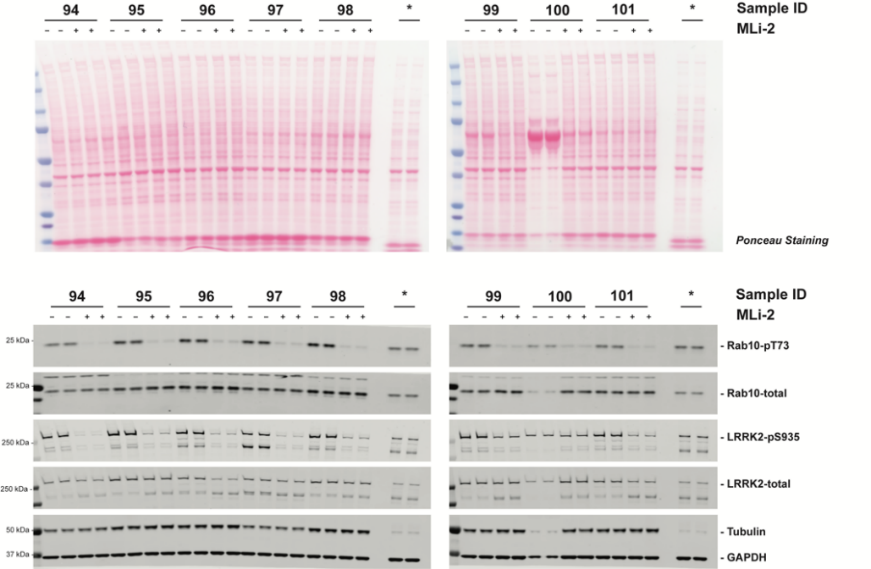

**Supplementary Figure 2: PBMC multiplexed quantitative immunoblot analysis for LRRK2 dependent Rab10 and Serine 935 LRRK2 phosphorylation.** PBMCs isolated from fresh peripheral blood were treated with either DMSO vehicle control or the specific LRRK2 kinase inhibitor MLi-2 at a concentration of 200nM for 30 minutes prior to cell lysis. 10 μg of whole cell extracts were then loaded in duplicates and subjected to quantitative immunoblot analysis with the indicated antibodies and the membranes developed using the Odyssey CLx scan Western Blot imaging system. pRab10 and total Rab10 protein as well as Serine 935 and total LRRK2 antibodies were multiplexed and the same internal standard was run on every gel to compare samples run on different gels (*). Corresponding Ponceau S staining of protein bands on the Western blot membranes is shown as well.

**Supplementary Tables**

**Supplemental Table 1.** Lymphoblastoid cell lines: genotypes, individual age at biopsy, sex, clinical status and ID numbers

**Cell line ID #	Age at biopsy	Genetic mutation	Clinical Status	Sex**

ND00011	75	None detected	Unaffected	F

ND02559	46	None detected	Unaffected	M

ND00312	72	None detected	Unaffected	M

ND01610	68	None detected	Affected	F

(wildtype for G2019S)

ND01979	75	None detected	Affected	F

(wildtype for G2019S)

ND00008	47	None detected	Affected	M

(wildtype for G2019S)

ND00318	72	None detected	Affected	M

(wildtype for G2019S)

**Supplemental Table 2. Demographics and clinical characterization of PD and healthy control subjects in archived PBMCs.**

Demographics	Control (n = 10)	PD (n = 53)

Age^a^ in years, (range)	63.9 (43,78)	65.3 (37,84)

Sex	6 F/ 4M	20 F/ 33 M

UPDRS part III (ON)	15.9

Age at diagnosis in years (range)	58.9 (35, 78)

Duration of disease in years (range)	5.9	(1, 19)

H & Y (mean)	2.0

^a^ Age at which the blood sample was obtained. No statistical difference was observed

between healthy controls and PD subjects (p = 0.7, student t-test).

**Supplemental Table 3. Demographics of healthy control subjects.**

Demographics	Control (n = 10)

Age^a^ in years, (± SEM)	62.3 (3.67)

Female %	50

European ancestry (%)	90

Ever Smoker (%)	20

Positive PD history^b^	20

^a^ Age at which the blood sample was obtained.

^b^ First degree relative with PD.

**Supplemental Table 4. Demographics and clinical characterization of PD and healthy control subjects in buffy-coat derived samples.**

Demographics	Control (n = 6)	PD (n = 14)

Age^a^ in years, (range)	64.8 (57, 69)	65.1 (53, 80)

Female %	33.3	14.2

European ancestry (%)	100	100

UPDRS part I	5.5	8.4

UPDRS part II	0.8	8.4

UPDRS part III^b^	2.83	27.1

Minutes since last dose	92.9

H & Y (mean)^c^	2.0

MOCA (mean)^c^	27.8	26.1

PDCRS^c^	105.2	96.6

Education in years	16.7	16.6

Smoker (%)	20	0

^a^ Age at which the blood sample was obtained. No statistical difference was observed

between healthy controls and PD subjects (p = 0.8, student t-test).

^b^ Part III UPDRS scores were obtained while PD patients were in the ON state (*78*).

^c^ Abbreviations: Montreal Cognitive Assessment (MOCA), Hoehn & Yahr stage (H&Y) and the Parkinson's Disease-Cognitive Rating Scale (PD-CRS).

**Supplemental Table 5. Demographics and clinical characterization of PD and healthy control subjects in buffy-coat derived samples obtained during a washout.**

Demographics	Control (n = 21)	PD (n = 16)

Age^a^ in years, (range)	65.2	67.5

Female %	61.9	18.8

European ancestry (%)	100	100

UPDRS part I	9.4

UPDRS part II	11.9

UPDRS part III^b^	25.7

UPDRS part III^c^	20.6

UPDRS total	42.6

^a^ Age at which the blood sample was obtained. No statistical difference was observed

between healthy controls and PD subjects (p = 0.8, student t-test).

^b^ Part III UPDRS scores were obtained while PD patients were in the OFF state.

^c^ Part III UPDRS scores were obtained while PD patients were in the ON state.

| **ID** | **Duke** | **Dundee** | **Sex** | **Group** | **Birth**  **Year** | **Age** | **Age at PD**  **diagnosis** | **PD**  **duration** | **UPDRS**  **Part III** | **H&Y** |
| --- | --- | --- | --- | --- | --- | --- | --- | --- | --- | --- |
| 1 | ✓ | ✓ | Female | Idiopathic PD | 1944 | 74 | 71 | 3 | 31 | 2 |
| 2 | ✓ | ✓ | Male | Idiopathic PD | 1941 | 77 | 73 | 4 | 42 | 2 |
| 3 | ✓ | ✓ | Male | LRRK2 PD | 1953 | 65 | 58 | 7 | 43 | 3 |
| 4 | ✓ | ✓ | Female | Healthy Control | 1953 | 66 |  |  |  |  |
| 5 | ✓ | ✓ | Female | Idiopathic PD | 1952 | 67 | 63 | 4 | 17 | 2 |
| 6 | ✓ | ✓ | Male | Idiopathic PD | 1949 | 70 | 63 | 7 | 37 | 3 |
| 7 | ✓ | ✓ | Male | Healthy Control | 1942 | 77 |  |  |  |  |
| 8 | x | ✓ | Male | LRRK2 NMC | 1979 | 40 |  |  |  |  |
| 9 | ✓ | ✓ | Female | LRRK2 NMC | 1977 | 42 |  |  |  |  |
| 10 | ✓ | ✓ | Female | Healthy Control | 1954 | 65 |  |  |  |  |
| 11 | ✓ | ✓ | Male | LRRK2 PD | 1943 | 76 | 72 | 4 | 25 | 2 |
| 12 | ✓ | ✓ | Female | LRRK2 PD | 1946 | 73 | 51 | 22 | 44 | 4 |
| 13 | ✓ | ✓ | Male | Healthy Control | 1951 | 68 |  |  |  |  |
| 14 | ✓ | ✓ | Male | LRRK2 NMC | 1963 | 56 |  |  |  |  |
| 15 | ✓ | ✓ | Male | Idiopathic PD | 1963 | 56 | 47 | 9 | 22 | 2 |
| 16 | ✓ | ✓ | Male | LRRK2 PD | 1953 | 66 | 38 | 28 | 33 | 2 |
| 17 | ✓ | ✓ | Female | Idiopathic PD | 1964 | 55 | 53 | 2 | 18 | 1 |
| 18 | ✓ | ✓ | Female | LRRK2 NMC | 1972 | 47 |  |  |  |  |
| 19 | ✓ | ✓ | Male | LRRK2 PD | 1946 | 73 | 58 | 15 | 42 | 2 |
| 20 | ✓ | ✓ | Female | Idiopathic PD | 1947 | 72 | 55 | 17 | 48 | 2 |
| 21 | ✓ | ✓ | Male | Idiopathic PD | 1958 | 61 | 58 | 3 | 22 | 2 |
| 22 | ✓ | ✓ | Male | Idiopathic PD | 1950 | 69 | 60 | 9 | 32 | 2 |
| 23 | ✓ | ✓ | Female | Healthy Control | 1955 | 64 |  |  |  |  |
| 24 | ✓ | ✓ | Male | Idiopathic PD | 1952 | 67 | 60 | 7 | 13 | 2 |
| 25 | ✓ | ✓ | Male | LRRK2 PD | 1950 | 69 | 59 | 10 | 38 | 2 |
| 26 | ✓ | ✓ | Male | LRRK2 PD | 1946 | 73 | 59 | 14 | 65 | 3 |
| 27 | ✓ | ✓ | Male | Idiopathic PD | 1939 | 80 | 75 | 5 | 26 | 2 |
| 28 | ✓ | ✓ | Male | LRRK2 PD | 1954 | 65 | 46 | 19 | 39 | 3 |
| 29 | ✓ | ✓ | Male | LRRK2 PD | 1935 | 84 | 79 | 5 | 37 | 2 |
| 30 | ✓ | ✓ | Male | Idiopathic PD | 1956 | 63 | 47 | 16 | 22 | 2 |
| 31 | ✓ | ✓ | Male | Idiopathic PD | 1947 | 72 | 53 | 19 | 39 | 2 |
| 32 | ✓ | ✓ | Male | Healthy Control | 1985 | 34 |  |  |  |  |
| 33 | ✓ | ✓ | Female | Idiopathic PD | 1956 | 63 | 49 | 14 | 40 | 2 |
| 34 | ✓ | ✓ | Male | LRRK2 PD | 1961 | 58 | 55 | 3 | 23 | 2 |
| 35 | ✓ | ✓ | Female | LRRK2 PD | 1957 | 62 | 59 | 3 | 22 | 2 |
| 36 | ✓ | ✓ | Female | LRRK2 PD | 1951 | 68 | 67 | 1 | 18 | 2 |
| 37 | ✓ | ✓ | Female | LRRK2 PD | 1934 | 85 | 77 | 8 | 30 | 2 |
| 38 | ✓ | ✓ | Male | LRRK2 PD | 1951 | 68 | 56 | 12 | 37 | 2 |
| 39 | ✓ | ✓ | Male | LRRK2 NMC | 1947 | 72 |  |  |  |  |
| 40 | ✓ | ✓ | Female | LRRK2 NMC | 1958 | 61 |  |  |  |  |
| 41 | ✓ | ✓ | Male | LRRK2 NMC | 1960 | 59 |  |  |  |  |
| 42 | x | ✓ | Female | LRRK2 NMC | 1962 | 58 |  |  |  |  |
| 43 | ✓ | ✓ | Female | LRRK2 NMC | 1954 | 66 |  |  |  |  |
| 44 | x | ✓ | Female | LRRK2 NMC | 1950 | 70 |  |  |  |  |
| 45 | ✓ | ✓ | Male | LRRK2 NMC | 1953 | 67 |  |  |  |  |
| 46 | ✓ | ✓ | Female | Healthy Control | 1950 | 68 |  |  |  |  |
| 47 | x | ✓ | Male | Healthy Control | 1979 | 39 |  |  |  |  |
| 48 | ✓ | ✓ | Female | Healthy Control | 1949 | 69 |  |  |  |  |
| 49 | x | ✓ | Female | Healthy Control | 1983 | 35 |  |  |  |  |
| 50 | ✓ | ✓ | Female | Idiopathic PD | 1942 | 76 | 71 | 5 | 16 | 2 |
| 51 | ✓ | ✓ | Male | Idiopathic PD | 1955 | 63 | 63 | 0 | 15 | 2 |
| 52 | ✓ | ✓ | Female | Healthy Control | 1955 | 63 |  |  |  |  |
| 53 | ✓ | ✓ | Male | Idiopathic PD | 1942 | 76 | 70 | 6 | 47 | 3 |
| 54 | x | ✓ | Female | Healthy Control | 1980 | 38 |  |  |  |  |
| 55 | ✓ | ✓ | Female | Idiopathic PD | 1946 | 72 | 51 | 21 | 8 | 1 |
| 56 | ✓ | ✓ | Female | Idiopathic PD | 1961 | 57 | 53 | 4 | 8 | 1 |
| 57 | ✓ | ✓ | Female | Healthy Control | 1955 | 63 |  |  |  |  |
| 58 | ✓ | ✓ | Female | LRRK2 NMC | 1985 | 33 |  |  |  |  |
| 59 | ✓ | ✓ | Male | Idiopathic PD | 1957 | 61 | 60 | 1 | 21 | 2 |
| 60 | ✓ | ✓ | Female | LRRK2 PD | 1976 | 42 | 33 | 9 | 26 | 3 |
| 61 | ✓ | ✓ | Female | LRRK2 NMC | 1948 | 70 |  |  |  |  |
| 62 | ✓ | ✓ | Male | Idiopathic PD | 1943 | 75 | 66 | 9 | 60 | 3 |
| 63 | ✓ | ✓ | Male | Healthy Control | 1980 | 38 |  |  |  |  |
| 64 | ✓ | ✓ | Female | Healthy Control | 1978 | 40 |  |  |  |  |
| 65 | ✓ | ✓ | Female | Idiopathic PD | 1951 | 67 | 58 | 9 | 6 | 2 |
| 66 | ✓ | ✓ | Male | Idiopathic PD | 1984 | 34 | 32 | 2 | 19 | 2 |
| 67 | ✓ | ✓ | Female | Healthy Control | 1953 | 65 |  |  |  |  |
| 68 | ✓ | ✓ | Female | LRRK2 NMC | 1987 | 32 |  |  |  |  |
| 69 | x | ✓ | Male | LRRK2 NMC | 1965 | 54 |  |  |  |  |
| 70 | ✓ | ✓ | Female | Healthy Control | 1976 | 42 |  |  |  |  |
| 71 | ✓ | ✓ | Male | Healthy Control | 1955 | 63 |  |  |  |  |
| 72 | x | ✓ | Male | Healthy Control | 1981 | 37 |  |  |  |  |
| 73 | ✓ | ✓ | Female | LRRK2 PD | 1962 | 57 | 53 | 4 | 11 | 1 |
| 74 | x | ✓ | Male | Healthy Control | 1980 | 39 |  |  |  |  |
| 75 | ✓ | ✓ | Male | Idiopathic PD | 1952 | 67 | 64 | 3 | 9 | 1 |
| 76 | ✓ | ✓ | Male | LRRK2 NMC | 1947 | 72 |  |  |  |  |
| 77 | ✓ | ✓ | Male | Idiopathic PD | 1947 | 72 | 64 | 8 | 35 | 3 |
| 78 | x | ✓ | Female | Healthy Control | 1942 | 77 |  |  |  |  |
| 79 | ✓ | ✓ | Male | LRRK2 NMC | 1958 | 61 |  |  |  |  |
| 80 | ✓ | ✓ | Male | LRRK2 NMC | 1959 | 60 |  |  |  |  |
| 81 | ✓ | ✓ | Female | LRRK2 PD | 1949 | 70 | 61 | 9 | 22 | 2 |
| 82 | ✓ | ✓ | Male | Idiopathic PD | 1938 | 81 | 72 | 9 | 45 | 2 |
| 83 | ✓ | ✓ | Male | Idiopathic PD | 1956 | 63 | 52 | 11 | 26 | 2 |
| 84 | ✓ | ✓ | Female | LRRK2 PD | 1950 | 69 | 64 | 5 | 9 | 2 |
| 85 | ✓ | ✓ | Male | Healthy Control | 1941 | 78 |  |  |  |  |
| 86 | ✓ | ✓ | Male | LRRK2 PD | 1952 | 67 | 58 | 9 | 5 | 2 |
| 87 | x | ✓ | Female | Healthy Control | 1952 | 67 |  |  |  |  |
| 88 | ✓ | ✓ | Male | LRRK2 PD | 1957 | 62 | 59 | 3 | 7 | 1 |
| 89 | ✓ | ✓ | Male | LRRK2 NMC | 1953 | 66 |  |  |  |  |
| 90 | ✓ | ✓ | Male | Idiopathic PD | 1954 | 66 | 56 | 10 | 45 | 2 |
| 91 | x | ✓ | Female | Healthy Control | 1946 | 74 |  |  |  |  |
| 92 | ✓ | ✓ | Female | LRRK2 PD | 1958 | 62 | 60 | 2 | 37 | 2 |
| 93 | ✓ | ✓ | Male | LRRK2 NMC | 1967 | 53 |  |  |  |  |
| 94 | ✓ | ✓ | Female | LRRK2 PD | 1957 | 63 | 44 | 19 | 18 | 2 |
| 95 | ✓ | ✓ | Male | LRRK2 PD | 1945 | 75 | 61 | 14 | 41 | 3 |
| 96 | x | ✓ | Female | Healthy Control | 1956 | 63 |  |  |  |  |
| 97 | x | ✓ | Female | LRRK2 NMC | 1962 | 57 |  |  |  |  |
| 98 | ✓ | ✓ | Male | LRRK2 PD | 1962 | 58 | 52 | 6 | 9 | 1 |
| 99 | ✓ | ✓ | Male | LRRK2 PD | 1951 | 69 | 59 | 10 | 24 | 2 |
| 100 | ✓ | ✓ | Female | LRRK2 PD | 1954 | 66 | 64 | 2 | 12 | 2 |
| 101 | ✓ | ✓ | Female | LRRK2 NMC | 1963 | 57 |  |  |  |  |
| 102 | ✓ | x | Female | Healthy Control | 1979 | 39 |  |  |  |  |
| 103 | ✓ | x | Male | Healthy Control | 1971 | 47 |  |  |  |  |
| 104 | ✓ | x | Female | LRRK2 PD | 1940 | 78 | 72 | 6 | 30 | 2 |
| 105 | ✓ | x | Male | Healthy Control | 1983 | 36 |  |  |  |  |
| 106 | ✓ | x | Male | Idiopathic PD | 1955 | 63 | 58 | 5 | 8 | 1 |
| 107 | ✓ | x | Male | LRRK2 PD | 1951 | 68 | 71 | 1 | 5 | 1 |
| 108 | ✓ | x | Female | Healthy Control | 1982 | 36 |  |  |  |  |
| 109 | ✓ | x | Male | Idiopathic PD | 1947 | 72 | 64 | 8 | 33 | 2 |
| 110 | ✓ | x | Male | Healthy Control | 1935 | 83 |  |  |  |  |
| 111 | ✓ | x | Male | Healthy Control | 1965 | 53 |  |  |  |  |

**Supplemental Table 6. Demographics and clinical characterization of the MJFF FBN cohort.** The MJFF FBN cohort consisted of 111 participants including healthy controls (n=31), idiopathic PD (n=30) and carriers of the *LRRK2* G2019S mutation (n=50) with PD (n=28) and non-manifesting carriers (n=22). A tick indicates whether a participant donated a blood sample for the mtDNA damage assay (Duke University, ‘Duke’), the LRRK2 dependent Rab10 phosphorylation assay (University of Dundee, ‘Dundee’) or both. X=did not donate blood for the respective study. Otherwise, biological sex, group, year of birth, age at study participation, age at PD diagnosis, PD duration in years, total UPDRS part III motor examination score and Hoehn & Yahr stage (H&Y) where applicable are described.

**Supplemental Table 7. Demographics of healthy control subjects.**

Demographics	Control (n = 20)

Age^a^ in years, (± SEM)	63.3 (1.81)

Female %	80

European ancestry (%)	80

Ever Smoker (%)	10

Positive PD history^b^	10

^a^ Age at which the blood sample was obtained.

^b^ First degree relative with PD.
